# State and trait characteristics of anterior insula time-varying functional connectivity

**DOI:** 10.1101/716720

**Authors:** Lorenzo Pasquini, Gianina Toller, Adam Staffaroni, Jesse A. Brown, Jersey Deng, Alex Lee, Katarzyna Kurcyus, Suzanne M. Shdo, Isabel Allen, Virginia E. Sturm, Yann Cobigo, Valentina Borghesani, Giovanni Battistella, Maria Luisa Gorno-Tempini, Katherine P. Rankin, Joel Kramer, Howard H. Rosen, Bruce L. Miller, William W. Seeley

**Author notes:** **Corresponding author:** William W. Seeley, MD. 675 Nelson Rising Lane 94158, San Francisco, California USA.

## Abstract

The human anterior insula (aINS) is a topographically organized brain region, in which ventral portions contribute to socio-emotional function through limbic and autonomic connections, whereas the dorsal aINS contributes to cognitive processes through frontal and parietal connections. Open questions remain, however, regarding how aINS connectivity varies over time. We implemented a novel approach combining seed-to-whole-brain sliding-window functional connectivity MRI and k-means clustering to assess time-varying functional connectivity of aINS subregions. We studied three independent large samples of healthy participants and longitudinal datasets to assess inter- and intra-subject stability, and related aINS time-varying functional connectivity profiles to dispositional empathy. We identified four robust aINS time-varying functional connectivity modes that displayed both “state” and “trait” characteristics: while modes featuring connectivity to sensory regions were modulated by eye closure, modes featuring connectivity to higher cognitive and emotional processing regions were stable over time and related to empathy measures.

## Introduction

The human anterior insula (aINS) is a functionally heterogeneous region implicated in functions ranging from interoceptive awareness and emotion processing to time perception and cognitive control (1). In humans, neuroimaging studies have begun to parcellate the aINS, based on its patterns of functional and structural connectivity. For example, task-free fMRI (tf-fMRI) studies, which measure the brain-wide correlation structure in slow (< 0.1 Hz), spontaneous blood oxygen level dependent (BOLD) signal fluctuations (2, 3), have shown that the ventral, agranular aINS is functionally connected to limbic and autonomic processing regions that include the pregenual anterior cingulate cortex, the amygdala, and subcortical structures such as the thalamus and periaqueductal gray (4–8). These regions make up the “salience network”, a large-scale distributed system that represents the homeostatic significance of prevailing stimuli and conditions (9–13). In contrast, the dorsal, dysgranular aINS is connected to a cingulo-opercular “task-control network” whose nodes include dorsolateral and opercular prefrontal, anterior midcingulate, and anterior parietal areas involved in cognitive control processes such as task-set initiation and maintenance (4, 7, 8, 14). Under task-free conditions, both aINS subregions, but perhaps especially the dorsal aINS (8, 15), show activity that is anticorrelated with the “default mode network”, a system including the posterior cingulate cortex, inferior parietal lobules, and precuneus (16). In addition to the dorsal-ventral axis, hemispheric lateralization of the aINS is proposed to help maintain bodily homeostasis by adjusting and balancing autonomic outflow based on bioenergetics demands (1, 10, 11). Recent evidence suggests that while the left-sided (dominant hemisphere) aINS controls parasympathetic tone, the homotopic right (non-dominant) aINS is more closely linked to sympathetic tone and responses (1, 10, 11). Structural and functional changes in aINS subregions have been reported in a variety of neuropsychiatric conditions, ranging from mood and anxiety disorders to schizophrenia, autism, and frontotemporal dementia (13, 17–20).

Standard tf-fMRI has helped to reveal the functional organization of the human aINS and other brain areas by providing a snapshot of functional connectivity as averaged across the duration of a scanning session. Although long-range structural connections are assumed to be relatively stable in the adult brain (21), coordinated functional activity is dynamic, with the brain continuously reshaping network configurations in response to prevailing conditions or task demands (22–24). Approaches that capture time-varying connectivity bring the potential to clarify how brain dynamics are organized and relate to function. We reasoned that this approach could help clarify how the aINS may change its network partners in response to salient internal and external stimuli and possibly contribute to the development of more granular and personalized fingerprints of brain function in health and disease. Such time-varying functional connectivity has been shown to relate to clinical outcomes (25–27); task performance (28); and behaviorally relevant measures of cognitive (29–31), emotional (32, 33), and attentional processing (34, 35). One of the few studies applying time-varying functional connectivity analyses to the insula parcellated it into posterior, middle, and dorsal and ventral anterior components, revealing partially overlapping time-varying connectivity profiles for ventral and dorsal aINS subregions. Ventral and dorsal profiles diverged based on distinct contributions from limbic/emotional processing and cingulo-opercular/cognitive regions mirroring findings from static tf-fMRI studies (15).

Despite this body of research, it remains unclear how distinct aINS subregions dynamically engage the salience and task-control networks over time. Moreover, it is largely unknown whether time-varying functional connectivity of the aINS — or any region for that matter — is modulated by externally-driven states or instead displays trait characteristics such as within-subject temporal stability and relationships to behavioral or dispositional measures. To gain a deeper understanding of these issues, we implemented a novel approach combining seed-to-whole-brain sliding-window functional connectivity and k-means clustering to derive time-varying functional connectivity profiles of aINS subregions across three large, independent samples of healthy participants, including eyes open vs. eyes closed conditions and longitudinal datasets. Across methods and samples, we found that aINS subregions display shared and distinct time-varying connectivity “modes” that bind together cognitive and/or emotional processing areas versus upstream sensory and motor cortices. Overall, the findings suggest that short-term temporal variability in aINS connectivity reflects both state and trait characteristics, revealing a path toward use of such data for assessing psychopharmacological treatment efficacy as well as long-term therapeutic disease modification in neuropsychiatric conditions.

## Materials and Methods

### Participants

All participants were healthy, cognitively normal adults, recruited from two different centers (Table 1, Supplementary Figure S1). Elderly participants were retrospectively selected from the Hillblom Aging Network, an extensively characterized longitudinal cohort assessed at the University of California, San Francisco (UCSF) Memory and Aging Center. Subjects were required to have a Clinical Dementia Rating Scale score (36) of 0 (range 0-3) and a Mini-Mental State Examination score (37) of 28 (range 0-30) or higher. Secondary inclusion criteria were based on availability of the Interpersonal Reactivity Index, a widely used questionnaire measuring emotional and cognitive empathy. Out of 224 elderly subjects with a tf-fMRI scan and at least one Interpersonal Reactivity Index available, a cross-sectional cohort of 121 elderly adults was selected based on availability of the Interpersonal Reactivity Index within three months of the tf-fMRI scan. A second sample was built based on availability of longitudinal tf-fMRI scans. Out of total 184 older adults with longitudinally assessed tf-fMRI, 68 were excluded because they overlapped with the cross-sectional sample (N = 121, described above) and 72 were excluded since they did not meet our selection criteria of being scanned twice within a 5-13 month interval. This procedure resulted in a final selection of 44 participants having longitudinal data and that did not overlap with the cross-sectional sample. All visits included neuropsychological testing and a neurologic exam in addition to a structural MRI and tf-fMRI scan. Exclusion criteria included a history of drug abuse, psychiatric or neurological conditions, and current use of psychoactive medications.

**Table 1.** Demographics and sample characteristics.

| Dataset | Cross-sectional | Longitudinal | Eyes closed/open |
| --- | --- | --- | --- |
| Number of subjects | 121 | 44 | 20 |
| Age in years, mean (s.d.) | 69.3 (3.6) | 72.9 (7.6) | 42.8 (6.4) |
| Sex (Female/Male) | 73/48 | 25/19 | 9/11 |
| Handedness (L/A/R) | 0/0/121 | 6/0/38 | 0/0/20 |
| Education in years, mean (s.d.) | 17.6 (2.0) | 17.5 (1.9) | 14.4 (2.6) |
| IRI-EC, mean (s.d.) | 27.4 (4.8) | NA | NA |
| IRI-PT, mean (s.d.) | 24.3 (5.7) | NA | NA |
| IRI-PD, mean (s.d.) | 13.1 (5.0) | NA | NA |
| IRI-FS, mean (s.d.) | 18.3 (5.1) | NA | NA |
| Interval between scanning dates in months, mean (s.d.) | NA | 9.2 (2.7) | 1.0 (0.2) |

A = ambidextrous; EC = empathic concern; FS = fantasy score; IRI = Interpersonal Reactivity Index; L = left; NA = not applicable; PD = personal distress; PT = perspective taking; R = right; s.d. = standard deviation

Twenty additional younger participants were recruited from a simultaneous FDG-PET/tf-fMRI study at the Klinikum rechts der Isar, Technische Universität München (38). Neuroimaging data was assessed twice within an interval of one month, during task-free conditions with either eyes closed or eyes open. Participants were randomly assigned to one of the two conditions, resulting in eight participants assessed with eyes closed at the first and with eyes open at the second scan, and 12 participants assessed with eyes open at the first and eyes closed at the second scan. Exclusion criteria included a history of psychiatric or neurological conditions, use of psychoactive medications, pregnancy, and renal failure.

For all samples, written informed consent was obtained from all involved participants and the study was approved by the institutional review board where the data was acquired (UCSF/Klinikum rechts der Isar).

### Empathy Assessment

The Interpersonal Reactivity Index (39) was completed by the participant’s informant within three months of tf-fMRI scan. This questionnaire is composed by four subscales, each one consisting of 7 questions that can comprehensively reach a maximum score of 35. While empathic concern and personal distress measure emotional aspects of empathy, perspective taking and the fantasy score are designed to assess cognitive aspects of empathy. The Interpersonal Reactivity Index was chosen since it has been widely used to assess different aspects of empathy in healthy and neuropsychiatric conditions (39, 40). For example, studies in frontotemporal dementia, a neurodegenerative diseases of socioemotional dysfunction, have shown that deficits in subscales of the Interpersonal Reactivity Index are linked to structural deterioration of the aINS and regions connected to the aINS such as the anterior cingulate and orbitofrontal cortices (40, 41).

### Neuroimaging data acquisitio

The cross-sectional and longitudinal cohorts from the Hillblom Aging Network were scanned at the UCSF Neuroscience Imaging Center on a Siemens Trio 3T scanner. A T1-weighted MP-RAGE structural scan was acquired with acquisition time=8 min 53 sec, sagittal orientation, a field of view of 160 × 240 × 256 mm with an isotropic voxel resolution of 1 mm^3^, TR=2300 ms, TE=2.98 ms, TI=900 ms, flip angle=9°. Task-free T2*-weighted echoplanar fMRI scans were acquired with an acquisition time=8 min 6 sec, axial orientation with interleaved ordering, field of view=230 × 230 × 129 mm, matrix size=92 × 92, effective voxel resolution=2.5 × 2.5 × 3.0 mm, TR=2000 ms, TE=27 ms, for a total of 240 volumes. During the 8-minute tf-fMRI acquisition protocol, participants were asked to close their eyes and concentrate on their breathing.

Data from the Klinikum rechts der Isar was acquired on an integrated Siemens Biograph scanner capable of simultaneously acquiring PET and MRI data (3T). FDG-PET activity and tf-fMRI was simultaneously measured during the initial 10 min immediately after bolus injection of the FDG tracer. Scanning was performed in a dimmed environment obtained by switching off all lights, including those in the scanner bore. Subjects were instructed to keep their eyes closed or open, to relax, to not think of anything in particular, and to not fall asleep. MRI data were acquired using the following scanning parameters: Task-free echoplanar fMRI scans: TR, 2.000 ms; TE, 30 ms/angle, 90°; 35 slices (gap, 0.6 mm) aligned to AC/PC covering the whole brain; sFOV, 192 mm; matrix size, 64 × 64; voxel size, 3.0 × 3.0 × 3.0 mm^3^ (each measurement consists of 300 acquisitions in interleaved mode with a total scan time of 10 min and 8 s); MP-RAGE: TR, 2.300 ms; TE, 2.98 ms; angle, 9°; 160 slices (gap, 0.5 mm) covering the whole brain; FOV, 256 mm; matrix size, 256 × 256; voxel size, 1.0 × 1.0 × 1.0 mm^3^; total length of 5 min and 3 s.

### Neuroimaging data preprocessing

Before preprocessing, all images were visually inspected for quality control. Images with excessive motion or image artifact were excluded. T1-weighted images underwent segmentation using SPM12 (http://www.fil.ion.ucl.ac.uk/spm/software/spm12/). For each tf-fMRI scan, the first five volumes were discarded. SPM12 and FSL (http://fsl.fmrib.ox.ac.uk/fsl) software were used for subsequent tf-fMRI preprocessing. The remaining volumes were slice-time corrected, realigned to the mean functional image and assessed for rotational and translational head motion. Volumes were next co-registered to the MP-RAGE image, then normalized to the standard MNI-152 healthy adult brain template using SPM segment, producing MNI-registered volumes with 2 mm^3^ isotropic resolution. These volumes were spatially smoothed with a 6 mm radius Gaussian kernel and temporally bandpass filtered in the 0.008-0.15 Hz frequency range using fslmaths. Nuisance parameters in the preprocessed data were estimated for the CSF using a mask in the central portion of the lateral ventricles and for the white matter using a highest probability cortical white matter mask as labeled in the FSL tissue prior mask. Additional nuisance parameters included the 3 translational and 3 rotational motion parameters, the temporal derivatives of the previous 8 terms (white matter/CSF/6 motion), and the squares of the previous 16 terms (42). Subjects were included only if they met all of the following criteria: no inter-frame head translations greater than 3 mm, no inter-frame head rotations greater than 3 degrees, and less than 24 motion spikes (defined as inter-frame head displacements > 1 mm), less than 10% of the total number of frames. Nuisance parameters were regressed out from the filtered data using fslmaths, and masked with a binarized, skullstripped MNI-152 brain mask. The WM/CSF/head movement denoised data was used for the subsequent time-varying functional connectivity analysis. Findings with global signal regression are not presented in the main body of the manuscript but were generated for the longitudinal dataset and are reported as sensitivity analysis in the Supplement.

### Time-varying functional connectivity analysis

For each individual, average blood oxygen level dependent signal time courses were extracted from the right and left ventral and dorsal aINS using four regions-of-interest from the Brainnetome Atlas (http://atlas.brainnetome.org/). In order to assess the impact of seed region selection, in the cross-sectional dataset activity time courses were also extracted from a language-relevant region centered on the left inferior frontal gyrus (IFG), using a region-of-interest defined in a previous study (43). Using in-house custom scripts based on Python (https://www.python.org/) and FSL, a sliding-window approach was implemented to generate time-varying seed-to-whole-brain connectivity maps (Figure 1A). The derived time series from the aINS and the entire tf-fMRI scan of each subject were divided into sliding-windows of 18 TRs (36 s) in steps of 1 TR creating 218 (275 for eyes closed/open) untapered, rectangular windows. At each window, linear regression was used to derive seed-to-whole-brain time-varying functional connectivity maps for each aINS seed (Figure 1B). A window size of 36 seconds was chosen based on previous research showing that window sizes between 30 and 60 seconds capture additional variations in functional connectivity not found in larger window sizes (24, 25, 44). Ideal sliding-window size has been explored by additional methodological work assessing the relationship between window length and cut-off frequencies, supporting the use of sliding-windows between lengths of 30–60 seconds for tf-fMRI data preprocessed using a low-pass filter set at 0.15 Hz (45). These studies are further supported by empirical findings showing that cognitive states can be discerned within such window lengths (46, 47). Nevertheless, in order to assess the impact of sliding-window length, control analyses were performed with window lengths of 72 TR (144 s). The resulting findings did not significantly differ from the reported findings with windows of 36 s (Supplementary Results and Figure S2); therefore 36 s windows were used throughout. The code used to derive time-varying functional connectivity maps is provided in the Appendix of the Supplement and in GitLab (https://gitlab.com/juglans/sliding-window-analysis/tree/master). The derived time-varying functional connectivity maps were finally standardized to z-scores (see Video 1 for left ventral aINS time-varying functional connectivity maps of a typical study participant), masked with a binarized gray matter mask, vectorized, and concatenated across subjects resulting in windows x voxels matrices for each aINS seed. K-means clustering was then applied to the concatenated window matrices using a k = 4 to produce four clusters representing four time-varying functional connectivity modes of aINS subregions present across the course of the functional scan (Figure 1C). The k-means algorithm used Euclidean distance, and an optimal solution was selected after 100 iterations and 10 replications. The optimal number of clusters, referred to henceforward as modes, was determined using elbow and silhouette plots and by performing additional clustering solutions with k = 3, 5, and 6 (Supplementary Results and Figure S3). With lower k solutions, the identified modes merged, resulting in information loss. Using higher k solutions resulted in the generation of redundant sub-modes, exemplified by Mode 4 of the left ventral aINS that would split in anterior and posterior centered components (Supplementary Figure S3B). A total of three clustering analyses per aINS subregion were independently performed on time-varying functional connectivity windows: one on the cross-sectional dataset; one on the longitudinal dataset; and one on the eyes closed/open dataset (see Supplementary Figure S1 for a schematized summary of k-means clustering analyses performed).

**Figure 1.**
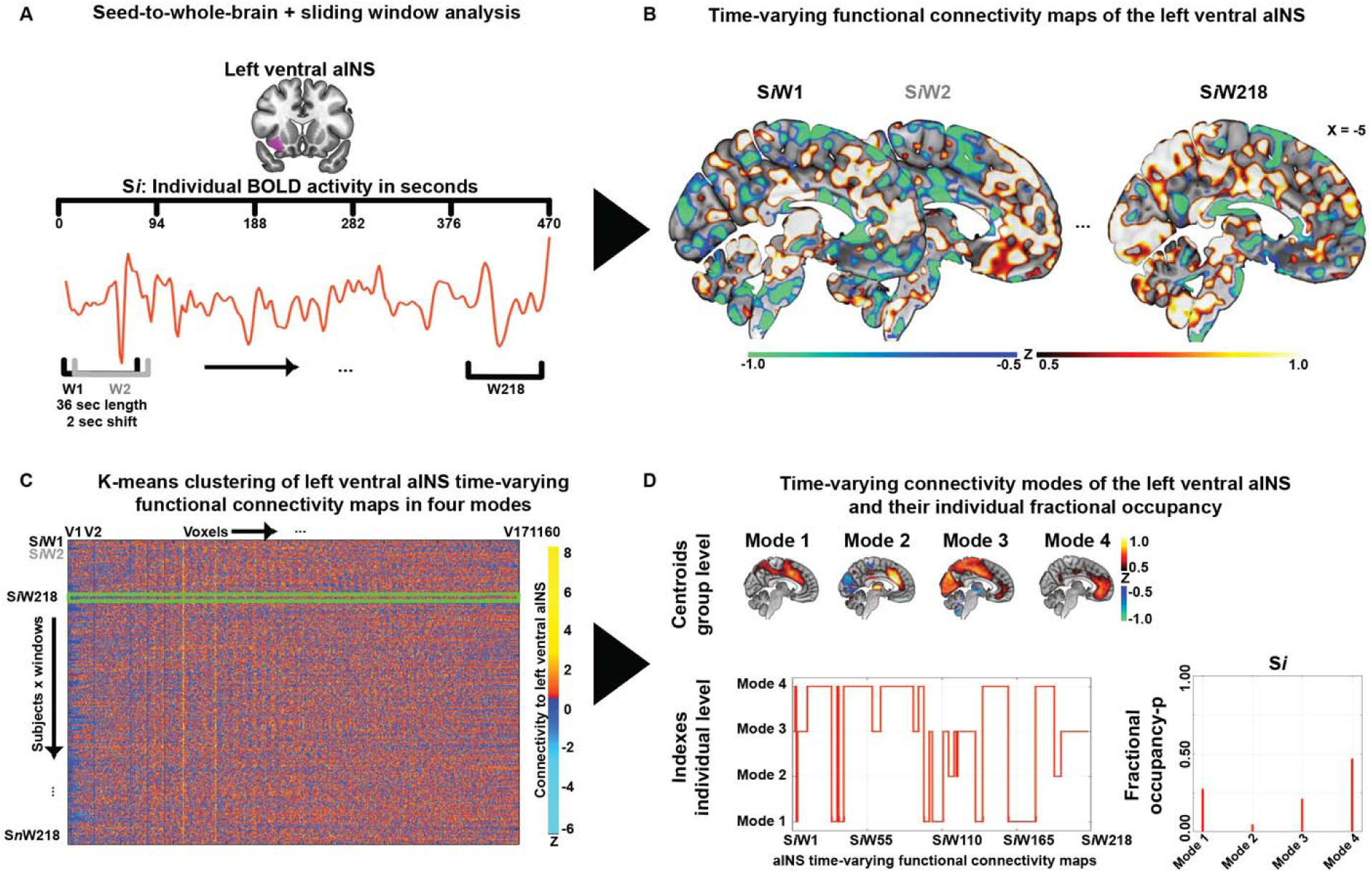
Time-varying functional connectivity pipeline. **(A)** For each individual S*i*, average BOLD activity time courses were extracted from subregions of the aINS, here the left ventral. **(B)** For each individual, 218 time-varying seed-to-whole-brain connectivity maps (S_*i*_W_1_-S_*i*_W_218_) were generated using a sliding-windows of 18 TRs (36 s) in steps of 1 TR. **(C)** The derived connectivity windows were vectorized and concatenated across subjects resulting in windows x voxels matrices for each aINS seed. K-means clustering was then applied to the concatenated window matrices using a value of k = 4 to produce four clusters representing four time-varying aINS modes present across the course of the functional scan. **(D)** Group averaged maps of the four modes were generated using the cluster-specific centroid maps generated through k-means (threshold at −0.5 > z > 0.5). The assignment of each window to a specific mode, was used to derive fractional occupancy profiles for each mode (i.e. the time spent by an aINS subregion in each mode), defined as the number of windows assigned to one mode divided through the total number of generated windows. aINS = anterior insula; BOLD = blood oxygen level dependent signal

Group-averaged maps of the four identified modes were generated, for each seed region, using the mode-specific centroid maps rendered using k-means clustering. The spatial similarity of modes derived from distinct aINS subregions and distinct groups was assessed by vectorizing the mode’s template maps and performing Pearson’s correlation analyses. To assess the distinct spatial contribution of major large-scale brain networks to time-varying connectivity modes, we used publicly available templates of major brain networks from a study investigating the functional network organization of the human brain (48)(https://www.jonathanpower.net/2011-neuron-bigbrain.html). Briefly, in this study subgraphs corresponding to major brain systems were derived from tf-fMRI data of > 300 healthy adults by retaining 2% of the strongest correlations. This procedure resulted in 12 binary templates spanning cognitive, primary sensory and subcortical systems: the salience, default, cingulo-opercular task-control, executive-control, ventral attention, dorsal attention, auditory, visual, ventral sensorimotor, dorsal sensorimotor, medial temporal lobe (“memory retrieval”) and subcortical networks (4, 48, 49) (Supplementary Figure S4). Subsequently, for each aINS subregion the averaged z-score value enclosed within these brain network templates was extracted from the mode-specific centroid maps generated in the cross-sectional dataset.

Further, the assignment of each window to a specific mode was used to derive: (i) the number of transitions from one mode to another; (ii) a fractional occupancy metric for each mode, defined as the number of windows assigned to that mode divided by the total number of windows; and (iii) how often aINS subregions simultaneously occupy distinct mode configurations in time. To identify meta-profiles of aINS time-varying functional connectivity, individual subject fractional occupancy profiles, defined as a vector of 16 elements summarizing the time fraction separately spent by individual aINS subregions on the four identified modes, were derived from the cross-sectional dataset. Fractional occupancy profiles were subsequently clustered using k-means with a clustering solution of k = 4 based on a silhouette analysis (using Euclidean distance, 100 iterations, and 10 replication). On the cross-sectional sample, static functional connectivity maps were also generated through voxel-wise regression analyses of each aINS seed’s time-course for the duration of the entire scan, following previous established methods (50, 51).

### Statistical analysis

Statistical analyses were carried out using R (https://www.r-project.org/) and Matlab-R2018 (https://www.mathworks.com/products/matlab.html). Pearson’s correlation was used to assess the spatial similarity of modes derived from different clustering analyses, using maps thresholded for z-values higher than 0.3 and lower than −0.3.

In the cross-sectional dataset, the Shannon diversity index was used to analyze whether aINS subregions of an individual subject transitioned repeatedly between distinct modes or showed instead a stereotypic behavior spending most of the time only in a specific aINS mode:

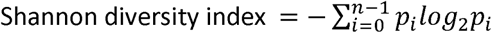

wherein *p* is the fractional occupancy of an aINS subregion in a specific time-varying functional connectivity mode. A Shannon diversity index of 1.4 indicates that all modes are equally occupied by the aINS, while the lower the Shannon diversity index, the higher the fractional occupancy of the aINS on one specific mode.

In the cross-sectional dataset, one-way ANOVAs were used to investigate whether aINS subregions differed, within subjects, in terms of mode-specific fractional occupancy, number of transitions, and Shannon diversity index. For descriptive purposes, averages and standard deviations derived over all aINS subregions and modes are reported. Additionally, for each meta-profile cluster identified in the cross-sectional dataset, one-way ANOVAs were used to assess differences in mode-specific fractional occupancy averaged across aINS subregions. The assignment of participant to four meta-profiles of time-varying aINS fractional occupancy was additionally used to test differences in aINS subregions’ static connectivity using ANOVA models and post-hoc t-tests implemented in SPM12 (height threshold p<0.005; extent threshold p<0.05 FWE corrected for multiple comparisons).

Finally, in the cross-sectional dataset, four multiple linear regression models were used to test the association of mode-specific fractional occupancies with subscales of the Interpersonal Reactivity Index (39). Subscales of the Interpersonal Reactivity Index were used as dependent variables, and each model contained the mode-specific fractional occupancies averaged across aINS subregions. Each model was corrected for age, sex, and sum frame-wise head displacement (p < 0.05 uncorrected for multiple comparisons). To assess specificity of the aINS results, four additional models were estimated, using subscales of the Interpersonal Reactivity Index as dependent variables and fractional occupancy of time-varying functional connectivity modes derived from the left IFG as predictors.

For the longitudinal datasets, cosine similarity was used to assess the similarity of fractional occupancy profiles across distinct scanning dates.

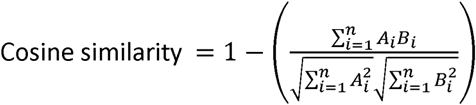

Where *A* and *B* are the vectorized fractional occupancy profiles of all aINS subregions at two different scanning dates. Cosine similarity varies between 1 and 0, with 1 indicating identical fractional occupancy profiles. Paired t-tests were performed to assess differences in fractional occupancy in the longitudinal and in the eyes closed/open datasets. All statistical findings are reported at p < 0.03, two-tailed, uncorrected for multiple comparisons except where specified otherwise.

## Results

### Bilateral ventral and dorsal aINS occupy four overlapping but distinct time-varying functional connectivity modes

By combining seed-to-whole-brain functional connectivity, sliding-window analysis, and k-means clustering, we found that aINS subregions adopt four overlapping but distinct large-scale configurations or “modes” of time-varying functional connectivity (Figure 2A). All modes were characterized by bilateral connectivity between the aINS and anterior cingulate cortices. Modes could be distinguished from each other, however, by specific patterns of connectivity to other brain regions (Figure 2A and Supplementary Table S1). In Mode 1, all four aINS subregions showed connectivity to the anterior midcingulate/pre-supplementary motor area, right frontal operculum, and dorsal parietal and dorsolateral prefrontal areas that together make up the task-control network. Mode 2 was characterized by negative connectivity to primary visual and sensorimotor areas and prominent connectivity to the ventral striatum and thalamus. Mode 3 showed inverted connectivity patterns to the same regions. Finally, Mode 4 showed connectivity patterns aligned with more ventral salience network regions, including pre- and sub-genual anterior cingulate and orbitofrontal cortices, with additional connectivity to the temporal poles for the right ventral aINS. All four modes were identified in each aINS subregion, as shown by the correlation matrix highlighting the spatial correspondence of equivalent modes (Figure 2B). Important distinctions were found, however, when comparing dorsal and ventral aINS connectivity patterns within modes. In Mode 1, the dorsal aINS showed prominent anti-correlations to the precuneus, angular gyrus, and medial prefrontal cortex regions that make up the default mode network (16). In Mode 4, the ventral aINS showed a more ventral connectivity pattern that encompassed subgenual anterior cingulate cortex and temporal poles, when compared to the dorsal aINS (Figure 2C, Supplementary Figure S6A). Modes derived from homologous left and right aINS regions differed from each other mainly with regard to the extent of ipsilateral connectivity to neighboring regions (Supplementary Figure S6B-C). Importantly, similar aINS time-varying functional connectivity modes were identified using longer sliding-window lengths (Supplementary Figure S2), lower and higher clustering solutions (Supplementary Figure S3), and data preprocessed using global signal regression (Supplementary Figure S7).

**Figure 2.**
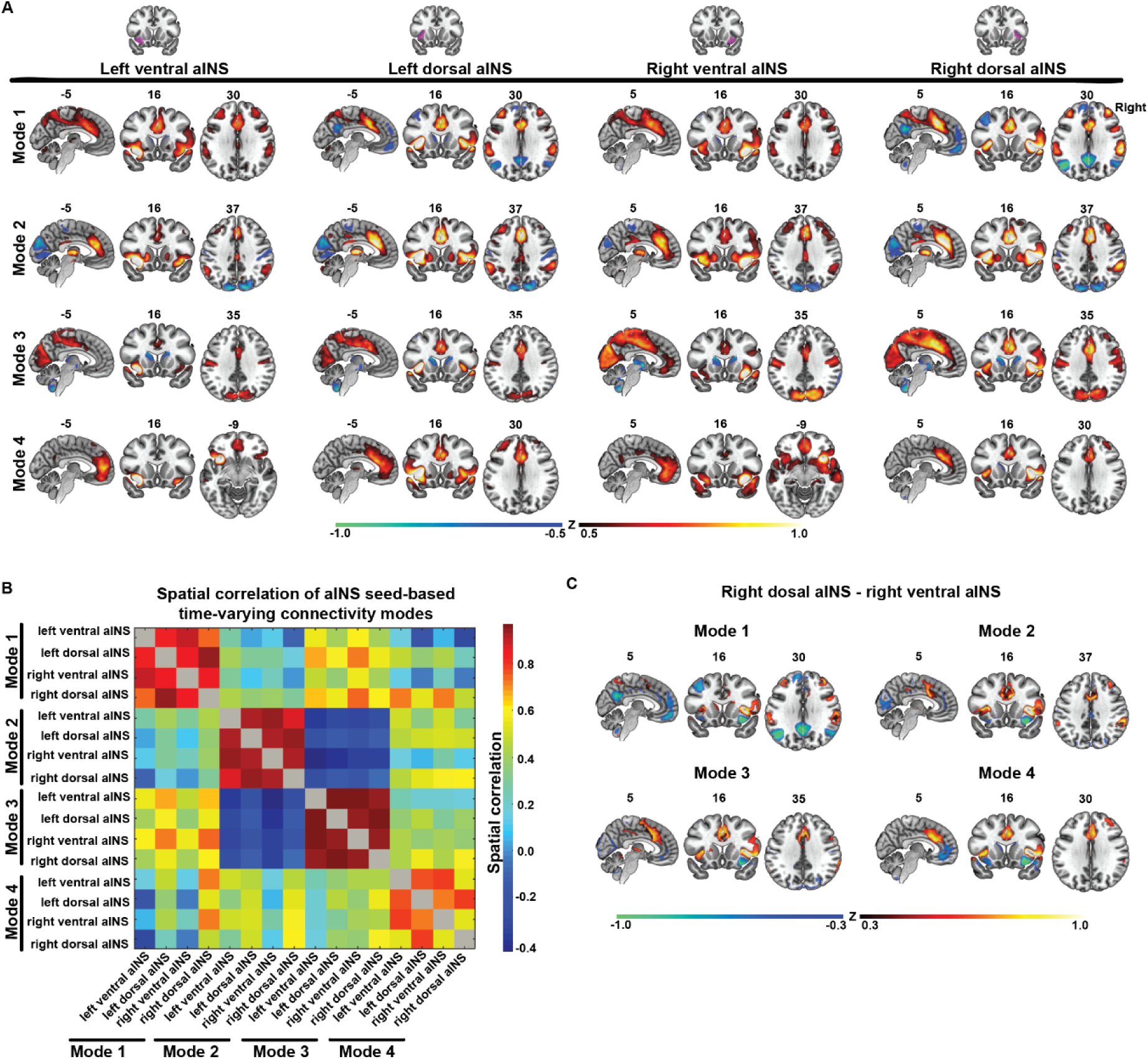
Time-varying functional connectivity modes of the aINS. **(A)** Spatial maps of the four time-varying functional connectivity modes identified across the four aINS subregions in the cross-sectional dataset. Threshold at −0.5 > z > 0.5, negative connectivity is depicted in blue, positive in red. The right side of the brain is shown on the right side of the image. **(B)** Correlation matrix reflecting the spatial similarity of modes identified in distinct aINS subregions. **(C)** Differences in time-varying connectivity between correspondent modes of the right dorsal and ventral aINS (threshold at −0.3 > z > 0.3, higher connectivity of the dorsal aINS is depicted in red, higher connectivity of the ventral aINS in blue).

### aINS subregions coherently transition between time-varying functional connectivity modes

Average fractional occupancy, i.e. the proportion of time spent by aINS subregions in the four time-varying functional connectivity modes ranged from 0.18 (left ventral aINS in Mode 2) to 0.30 (right ventral aINS in Mode 1), with an overall average of 0.25 +/-0.20 (Figure 3A). Importantly, no significant differences in mode-specific fractional occupancy were found across the four aINS subregions (Mode 1: F = 0.2, p = 0.82; Mode 2: F = 0.7, p = 0.56; Mode 3: F = 0.4, p = 0.77; Mode 4: F = 0.5, p = 0.71). In only a few participants did aINS subregions spend disproportionate time in only one mode. The number of transitions was also comparable across aINS subregions (F = 0.4, p = 0.759; Figure 3B), with an overall average of 18.0 +/-6.6 transitions over the task-free acquisition. In particular, our data shows that there were no direct transitions between the primary sensory anticorrelated Mode 2 and the primary sensory correlated Mode 3 (Supplementary Figure S8), suggesting that aINS subregions need to transition to modes rooted in cognitive networks before occupying primary sensory-centered modes characterized by opposing connectivity patterns. The Shannon diversity index was used to quantify the diversity of fractional occupancy and to compare this diversity across subregions. On average, aINS subregions showed high Shannon diversity, suggesting that each subregion moved between modes in most subjects, rather than stacking on a single mode. The overall average Shannon diversity index of fractional occupancy across aINS subregions was 1.0 +/-0.2 and did not significantly differ between subregions (F = 0.8; p = 0.503; Figure 3C). Subject-level fractional occupancy profiles are illustrated in Figure 3D, which shows two subjects with contrasting signatures. We finally sought to assess whether aINS subregions coherently occupy the same modes at the same time, or for instance, how often the right ventral and dorsal aINS are simultaneously connected to Mode 1. For each subject and each aINS subregion, we assessed how often possible mode configurations were jointly occupied in time. This analysis revealed a tendency across aINS subregions to occupy the same modes in time (Figure 3E). On average, the right ventral and dorsal aINS occupied the same modes 50% of the time, the left ventral and dorsal aINS 53% of the time, the right and left ventral aINS in 51% of the time, and the right and left dorsal aINS 61% of the time. These findings suggest an overall tendency for aINS subregions to cohesively engage the same modes over time under task-free scanning conditions.

**Figure 3.**
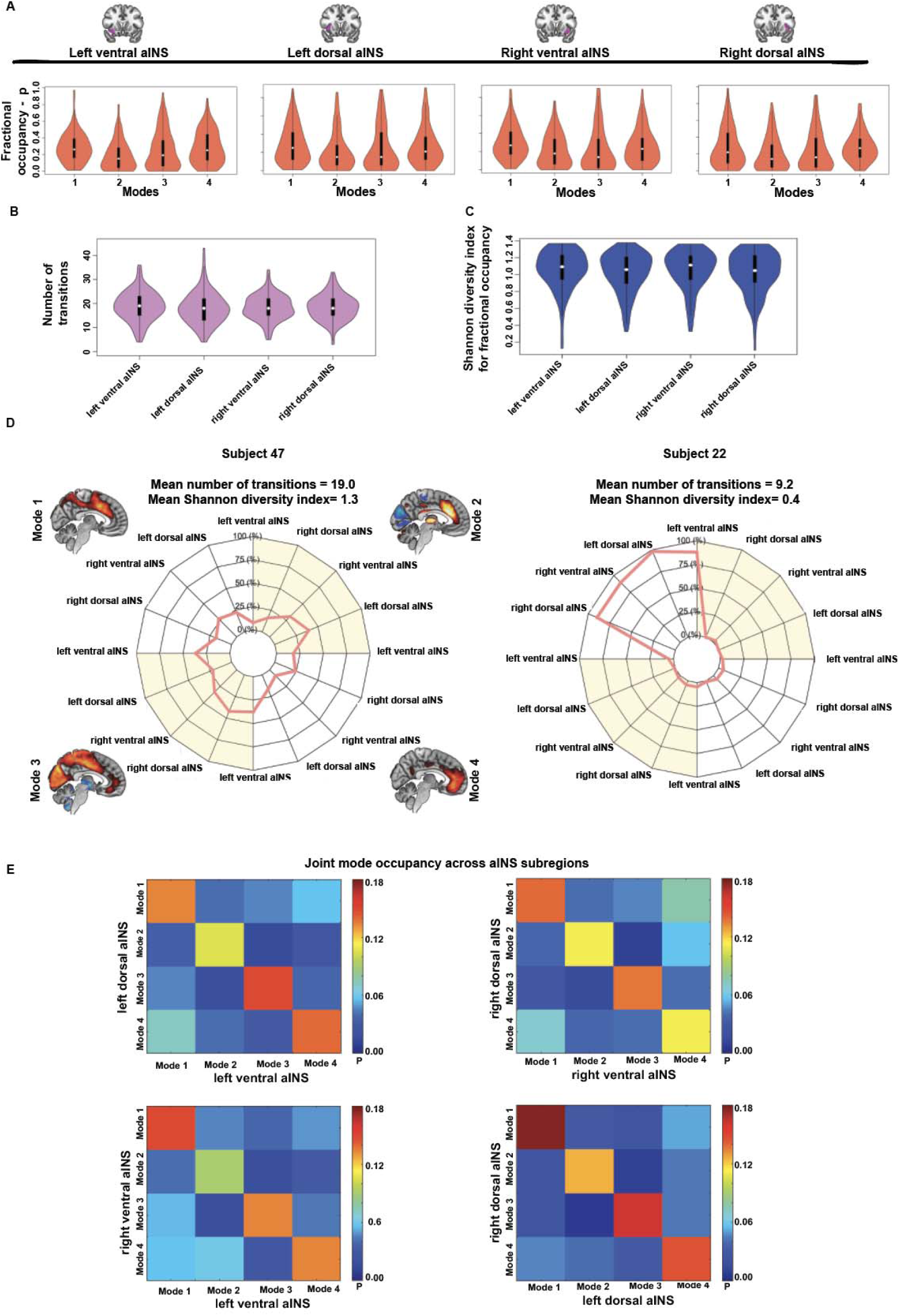
Temporal characteristics of the identified aINS time varying functional connectivity modes. **(A)** Violin plots reflecting group fractional occupancy in the four identified time-varying functional connectivity modes across the four aINS subregions. **(B)** Violin plots representing the number of transitions between time-varying functional connectivity modes across the four aINS subregions. **(C)** Violin plots representing the diversity of fractional occupancy profiles – measured using the Shannon diversity index – across aINS subregions. This measure reflects whether an aINS subregion tends to spend similar amounts of time in different modes or spends most of its time in only one mode. **(D)** Polar plots schematizing the fractional occupancies of aINS subregions into the four time-varying functional connectivity modes. On the left side, the fractional occupancy profile of Subject 47 transitions between several modes, and is characterized by a high number of transitions and by high Shannon diversity index averaged across aINS subregions. On the right side, we can appreciate the highly stereotyped fractional occupancy profile of Subject 22, with aINS subregions spending most of their time in a single mode. Regardless of the subregion, aINS subregions in this participant tend to spend time only in Mode 1, showing hence a low number of transitions and low Shannon diversity index averaged across aINS subregions. **(E)** Heat maps reflecting how specific modes are jointly occupied at the same time by distinct aINS subregions. Scale bar reflects the average group probability that distinct aINS subregions occupy certain mode configurations.

### Individual aINS connectivity mode occupancy profiles are reproducible across samples and within subjects over time

To assess the generalizability and reproducibility of our findings, we next turned to a longitudinal healthy aging dataset, which consisted of 44 participants who were scanned twice over an interval between 5-13 months. Clustering of time-varying connectivity windows identified modes that strongly resembled the four modes derived from the larger (and non-overlapping) cross-sectional dataset (Figure 4A-D and Supplementary Figure S5 panels A, C and E). Average fractional occupancies of aINS subregions in the four modes were comparable with the cross-sectional sample (comparing Figure 3A to 4A-D). Across aINS subregions and modes, paired t-tests revealed no significant differences in fractional occupancy when comparing the first scanning date with the second (p < 0.05 uncorrected for multiple comparisons, for details on paired t-test statistics see Supplementary Table S2). Within subjects, cosine similarity analysis showed that most participants had relatively stable fractional occupancy profiles over time (mean cosine similarity = 0.70 +/-0.19 s.d.), as further supported by the inverted bell-shaped distribution of cosine similarity (Figure 5A). Stable fractional occupancies are exemplified in Subjects 10 (first scanning date in red; second scanning date in blue; cosine similarity = 0.98) and 41 (cosine similarity = 0.92) (Figure 5B-C). A few participants, however, showed very different fractional occupancy profiles when comparing the two scanning dates, as exemplified by Subjects 21 (cosine similarity = 0.46) and 26 (cosine similarity = 0.24) (Figure 5D-E). Overall, however, the temporal stability of the individual profiles suggests a major contribution from trait-level factors.

**Figure 4.**
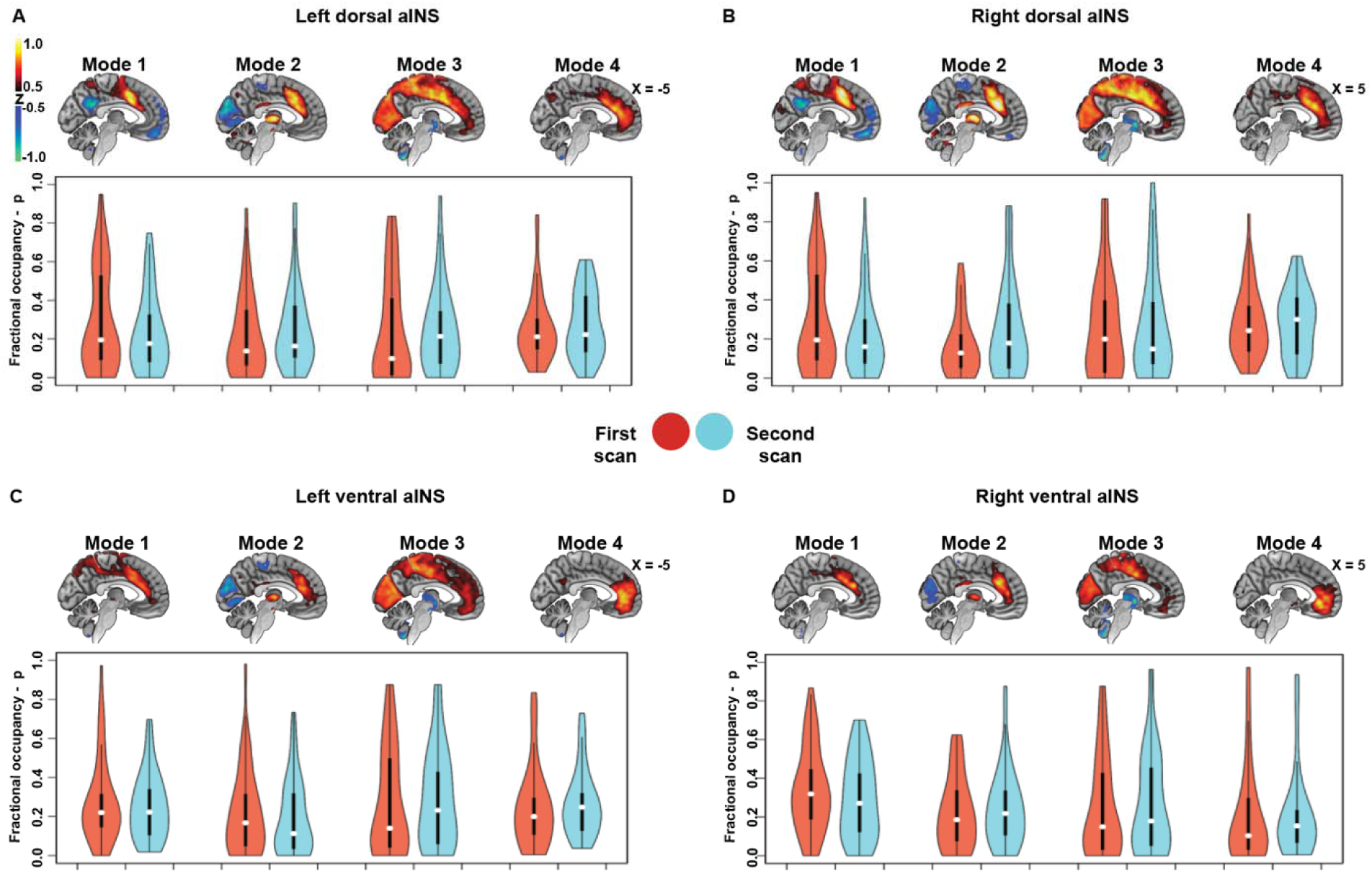
Time-varying functional connectivity modes across time. In the longitudinal data sample, spatial patterns of time-varying functional connectivity modes where identified across the four aINS subregions bearing high similarity to the modes identified in the cross-sectional sample (maps averaged across both time points, sagittal plane only panels **A-D**). Fractional occupancy in the identified modes did not significantly differ between the first (red violin plots) and second scanning dates (blue violin plots) ∼9 months apart (paired t-test; p < 0.05 uncorrected for multiple comparisons).

**Figure 5.**
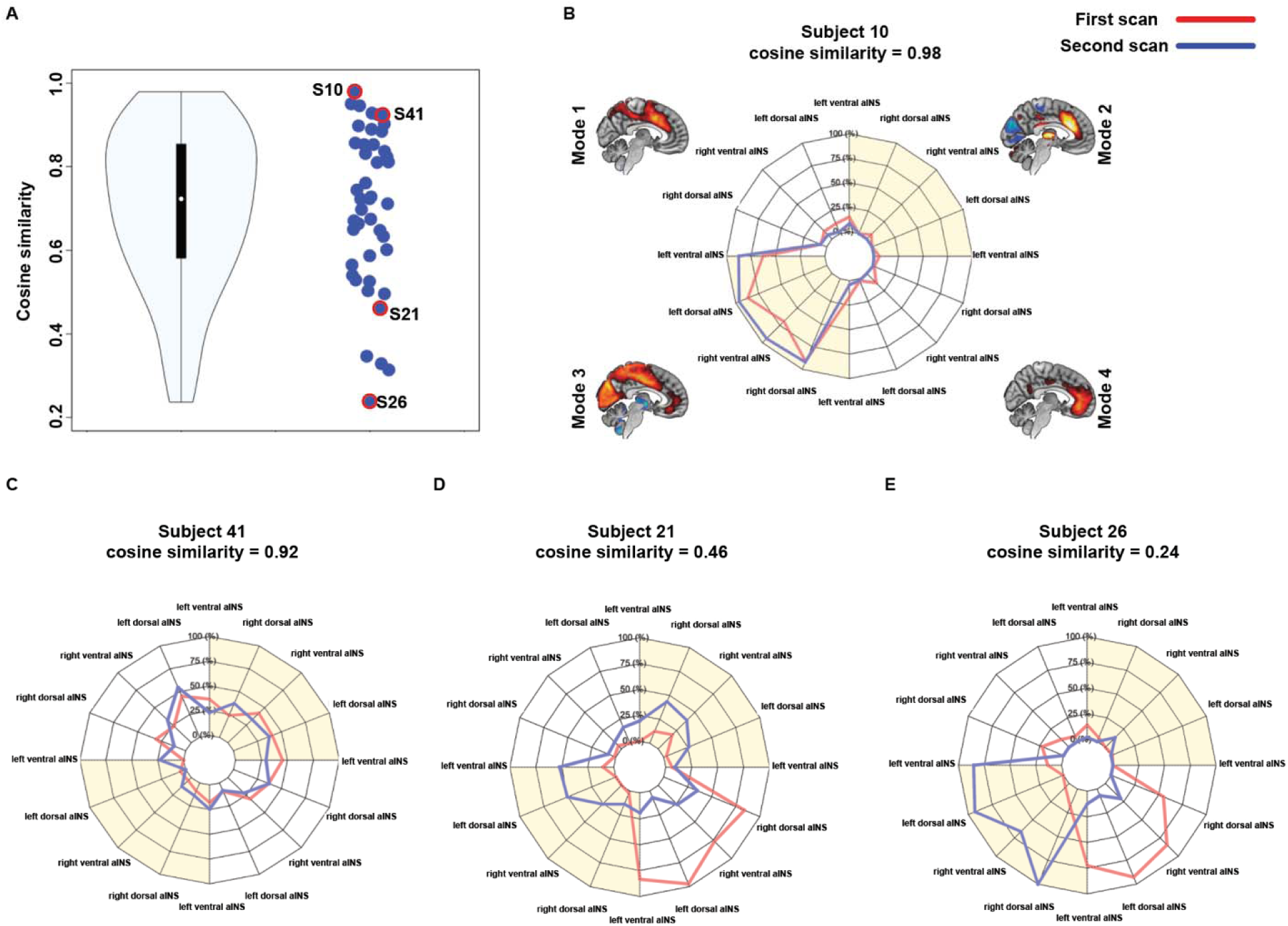
Individual fractional occupancy of aINS subregions across time. **(A)** Violin plot representing the similarity of aINS fractional occupancy profiles across scanning dates assessed using cosine similarity. Subjects 10, 41, 21 and 26 are highlighted with red circles in the individual data plot. The individual fractional occupancy profiles of these subjects and correspondent cosine similarity values are further schematized in panels **B-E**, where the fractional occupancies are shown at the first (red) and second (blue) scanning dates. aINS fractional occupancy in Subjects 10 and 41 did not substantially differ between scanning dates (note the overlap in the polar plots and the high cosine similarity index), while Subjects 21 and 26 showed very different aINS fractional occupancy profiles when assessed at different time points (note the little overlap in the polar plot and the low cosine similarity index).

### Eye opening decreases the time aINS spends anticorrelated with visual cortices

Independently performed clustering of time-varying connectivity data from the eyes closed/open dataset revealed similar aINS modes to those identified in the cross-sectional and longitudinal aging datasets (Figure 6A-B and Supplementary Figure S5 panels B, D and E). For all subregions except the right ventral aINS, participants showed higher fractional occupancies in the anti-correlated primary sensory/motor Mode 2 with eyes closed and higher fractional occupancies in the task-control Mode 1 with eyes open (p < 0.05 uncorrected, for details on paired t-test statistics see Supplementary Table S2).

**Figure 6.**
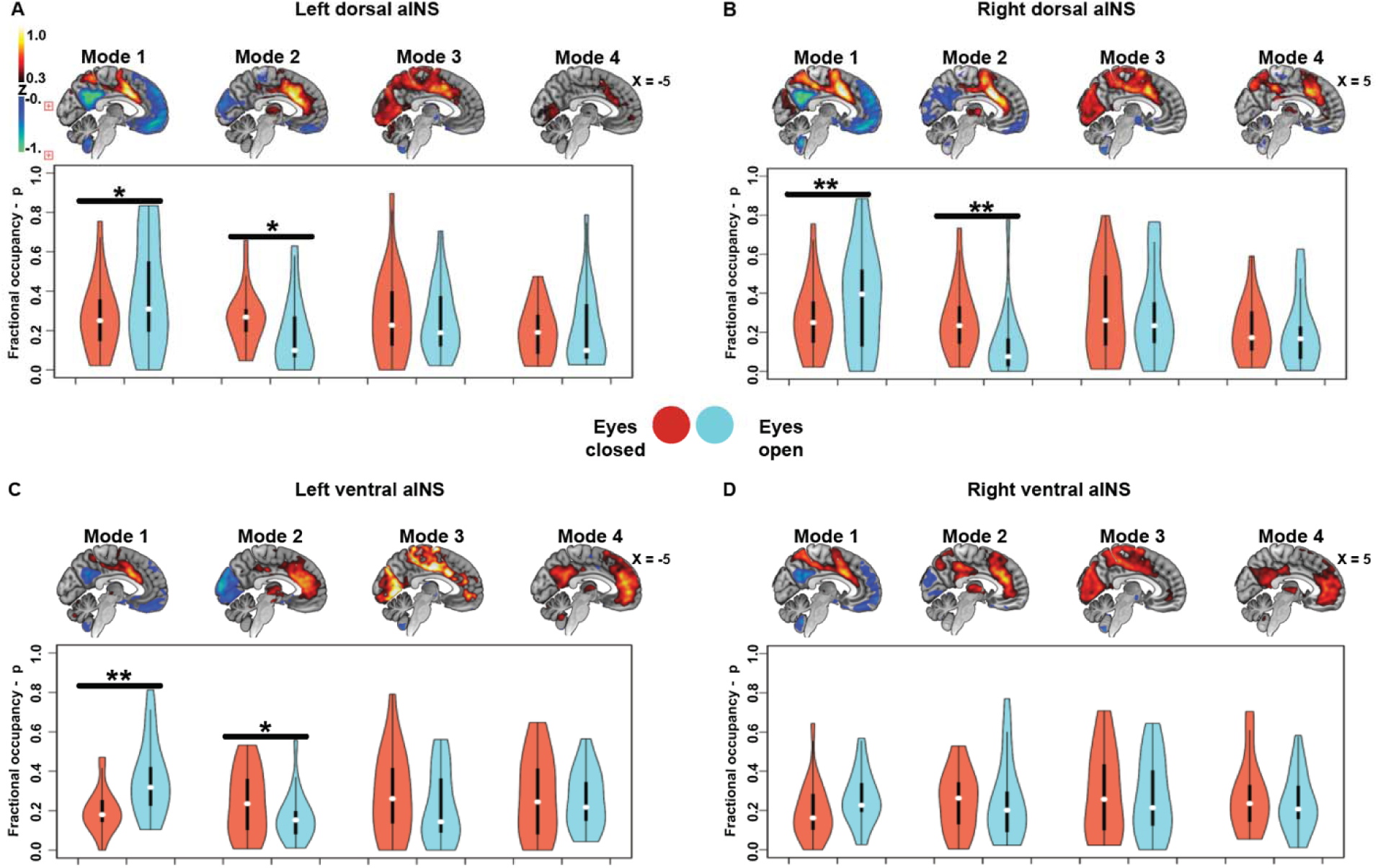
Time-varying functional connectivity modes during scanning with eyes closed versus eyes open. In the eyes closed/open dataset, spatial patterns of time-varying functional connectivity modes where identified across the left dorsal, right dorsal, left ventral, and right ventral aINS bearing high similarity to the modes identified in the cross-sectional sample (maps averaged across eyes closed/open conditions, sagittal plane only panels **A-D**). With exception of the right ventral aINS, fractional occupancy in the visually anticorrelated Mode 2 was significantly higher during eyes closed (red violin plots) than during eyes open scanning (blue violin plots). On the other hand, in the eyes open condition participants spent significantly more time in the task-control Mode 1 (paired t-test; p < 0.05 uncorrected for multiple comparisons). *p < 0.05; **p < 0.005

### Time-varying functional connectivity meta-profiles show that aINS subregions cohesively occupy the same modes

Individual subjects were clustered into meta-profiles (see Methods “*Time-varying functional connectivity analysis*”), based on the tendency of their aINS subregions to occupy specific time-varying functional connectivity modes. Individual fractional occupancy profiles of the cross-sectional sample (n = 121) were clustered into four clusters using k-means. Silhouette plots were used to choose the ideal number of clusters, k = 4 (Supplementary Figure S9). This analysis revealed four meta-profiles, characterized by preferred fractional occupancy in one of the four time-varying functional connectivity modes (Figure 7A). Meta-profile 1 (in green, 26 subjects) was characterized by a tendency to spend time in the task-control Mode 1 (see Subject 22, Figure 7A); Meta-profile 2 (in blue, 31 subjects) was characterized by a tendency to spend time in the sensory anticorrelated Mode 2 (see Subject 79, Figure 7A); Meta-profile 3 (in violet, 44 subjects) was characterized by a tendency to spend time in the sensory connected Mode 3 (see Subject 80, Figure 7A); while Meta-profile 4 (in red, 20 subjects) was characterized by a tendency to spend time in the salience network Mode 4 (see Subject 88, Figure 7A). To evaluate statistical differences in mode occupancy across meta-profiles, we averaged the time spent in each mode across aINS subregions and used one-way ANOVAs to test for significant group differences in mode-specific fractional occupancy across the distinct aINS meta-profiles (Meta-profiles 1, F = 50.1, p < 0.0001; Meta-profile 2, F = 39.8, p < 0.0001; Meta-profile 3, F = 79.6, p < 0.0001; Meta-profile 4, F = 46.1, p < 0.0001, see also Figure 7B). These differences were further statistically assessed via post-hoc t-tests. In summary, these analyses revealed a preference for aINS subregions to collectively spend similar fractions of time in a given mode, whereas distinct modes were primarily occupied by subjects in different meta-profiles.

**Figure 7.**
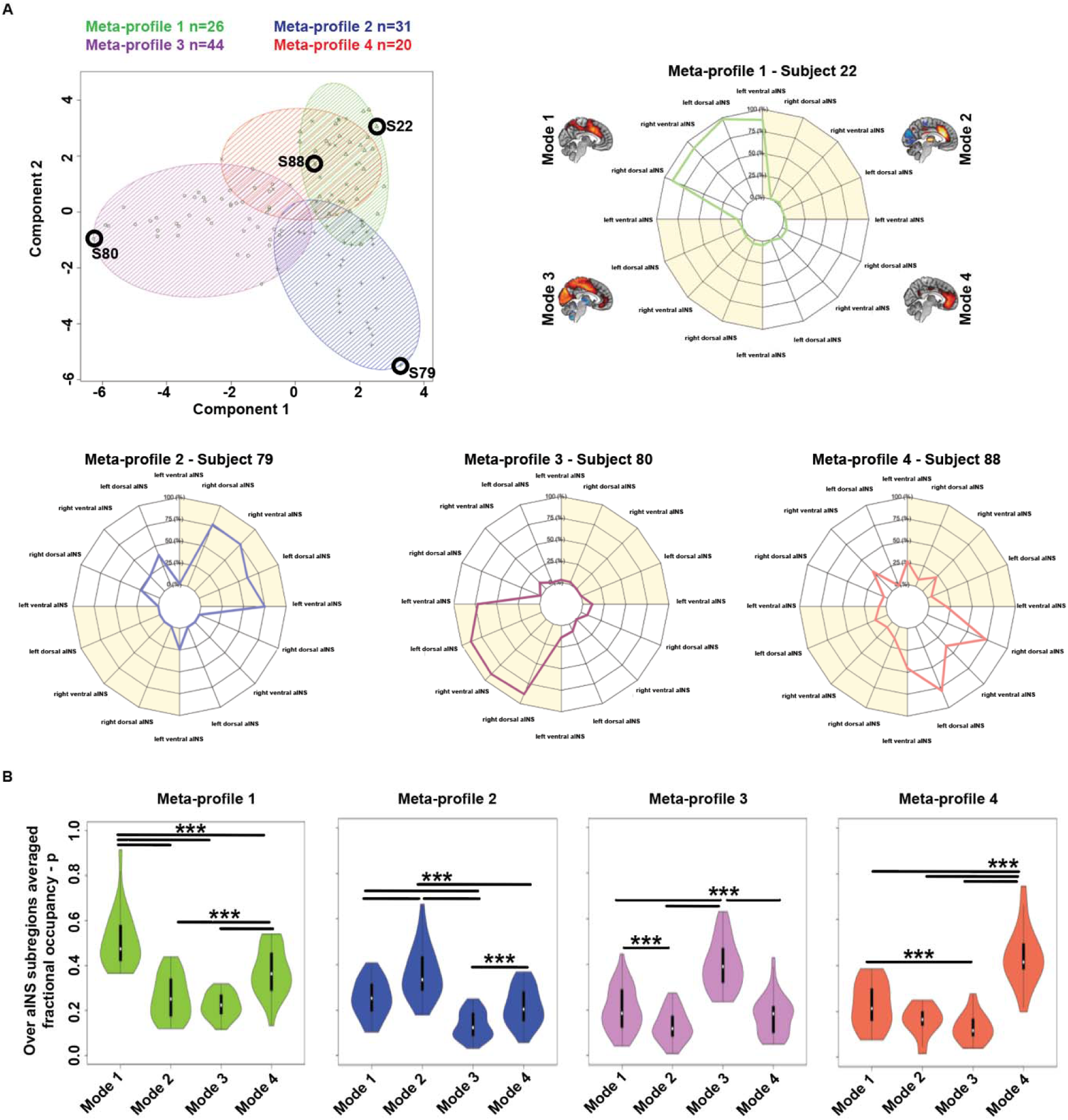
Meta-profiles of time-varying dynamic connectivity modes. **(A)** Individual fractional occupancy profiles of aINS subregions were clustered into four clusters, reflecting four meta-profiles of time-varying aINS functional connectivity. 26 subjects were clustered into Meta-profile 1 (triangles, green), 31 into Meta-profile 2 (+ sign, blue), 44 into Meta-profile 3 (circles, violet), and 20 into Meta-profile 4 (x’s, red). Panels show individual fractional occupancy profiles representative of the identified meta-profiles (see black circles in the scatterplots). Across aINS subregions, Subject 22 tended to spend time in Mode 1, Subject 79 in Mode 2, Subject 80 in Mode 3, and Subject 88 in Mode 4. **(B)** When averaged across aINS subregions, one-way ANOVAs and related post-hoc t-tests reveal that the aINS of participants clustered in Meta-profile 1 showed significantly higher fractional occupancy in Mode 1. Similarly, the aINS of participants clustered on Meta-profiles 2, 3, or 4, spent more time on Modes 2, 3, or 4, respectively.

Next, we asked whether the constituents of a given meta-profile would show group-level differences in terms of their static functional connectivity patterns. We therefore compared groups based on static functional connectivity of aINS subregions across the four meta-profiles described above. This analysis revealed that participants in Meta-profile 1 showed prominent static anticorrelation of the dorsal aINS to the default mode network when compared to participants clustered on other meta-profiles. Participants clustered on Meta-profile 2 and 3 showed marked visual and sensorimotor cortex static hypoconnectivity and hyperconnectivity, across all aINS subregions, when compared to participants of other meta-profiles; while participants in Meta-profile 4 showed prominent static connectivity of all aINS subregions to more ventral frontal regions (Supplementary Figure S10). In summary, key components of the time-varying connectivity modes were reflected in static connectivity differences between meta-profile-based groups.

### Comparison of aINS and left IFG time-varying functional connectivity

To control for the location of the seed, we derived time-varying functional connectivity modes by seeding the left IFG. Time-varying functional connectivity modes of the left IFG shared some common themes with modes identified using aINS subregions (Supplementary Figure S11A). The four identified modes showed similar connectivity to regions of the language network, such as the left IFG, pre-supplementary motor area, and superior parietal lobule (43, 52). This contrasts with the modes identified in aINS subregions that showed overlapping time-varying functional connectivity to typical task-control and salience network regions (Supplementary Figure S11B). Modes derived from both the aINS and left IFG, however, showed similar patterns of default mode network anticorrelation (Mode 1), primary sensory anticorrelation (Mode 2), and primary sensory hypercorrelation (Mode 3). A fourth mode was identified using the left IFG, showing time-varying functional connectivity to regions of the dorsal attention network (49) (Mode 4).

### Time spent by the aINS in the task-control and salience networks correlates with metrics related to dispositional empathy

A key question of this work concerns whether individual differences in aINS time-varying connectivity relate to differences in aINS-associated traits and functions. Among the many tasks that activate and depend on the anterior insula, empathy is one of the best documented (1, 18). The Interpersonal Reactivity Index is an informant-based questionnaire widely used to assess distinct aspects of empathy (39). Emotional aspects of empathy are covered by the empathic concern and personal distress measures, with the first assessing another-centered emotional response, while the second reflects general anxiety and self-oriented emotional reactivity. Perspective taking and fantasy scores measure cognitive aspects of empathy, with the first assessing the tendency to spontaneously imagine the cognitive perspective of another person, while the second measures the tendency to project oneself into the experiences of fictional characters. Mode-specific fractional occupancies were averaged across aINS subregions and used as regressors in four models using subscales of the Interpersonal Reactivity Index as dependent variables. These multiple linear regression models were corrected for age, sex, and sum frame-wise head displacement and identified two significant positive associations with time-varying connectivity metrics. First, fractional occupancy in the task-control Mode 1 predicted higher levels of personal distress, reflecting self-oriented feelings of personal anxiety and unease in tense interpersonal settings (β = 7.1; p < 0.05 uncorrected for multiple comparisons; Supplementary Table S3). Second, fractional occupancy in the salience Mode 4 correlated with higher scores on the fantasy subscale, reflecting greater ability to transpose one’s self imaginatively into the feelings and actions of fictitious characters in books, movies, and plays (β = 9.2; p < 0.05 uncorrected for multiple comparisons; Supplementary Table S3). These results remained significant when removing outliers (Mode 1 and personal distress β = 6.9, p < 0.05 uncorrected for multiple comparisons; Mode 4 and fantasy score β = 9.2, p < 0.05 uncorrected for multiple comparisons). No significant associations were found between fractional occupancy in any mode and the empathic concern or perspective taking subscales. As hypothesized, fractional occupancy of left IFG time-varying connectivity modes was not significantly associated with any measure of dispositional empathy (p < 0.05 uncorrected for multiple comparisons, see Supplementary Table S3).

## Discussion

To date, most efforts to relate large-scale networks to individual differences have relied on metrics that capture the topology or strength of node-to-node static functional (or “intrinsic”) connections. Here, we developed a novel approach combining seed-to-whole-brain functional connectivity, sliding-window analysis, and k-means clustering on tf-fMRI data to identify distinct time-varying functional connectivity modes of the aINS, a brain region critical for many human socio-emotional functions (1, 18). These robust modes were identified across methods and samples, showing both “state” and “trait” characteristics. In particular, while aINS modes related to sensory processing were modulated by visual input (eyes closed vs. open conditions during scanning), modes featuring connectivity to cognitive and emotion processing regions were stable over time and related to measures of dispositional empathy.

### Partially overlapping but distinct time-varying aINS functional connectivity modes

Our pipeline reliably identified four time-varying functional connectivity modes of the aINS across different subregions, methods, and samples. These modes were characterized by common connectivity of aINS subregions to the contralateral insula and to the anterior cingulate cortex, but could be differentiated from each other based on recruitment of other brain regions. Mode 1 was characterized by a more dorsal connectivity pattern that resembles previous characterizations of a cingulo-opercular task-control network (4, 14, 53) involving the bilateral aINS and anterior cingulate cortex, with additional connectivity to dorsomedial parietal and dorsolateral frontal areas. Mode 2 was characterized by negative connectivity to primary sensory areas and increased connectivity to the thalamus while Mode 3 showed an inverted connectivity pattern to these regions. Finally, Mode 4 was characterized by fronto-insular connectivity to more rostral and ventral portions of the anterior cingulate cortex, resembling the salience network (4, 8), with additional connectivity to the temporal pole and medial orbitofrontal cortex, components of the “semantic-appraisal network” (18, 54). These findings are consistent with those of a previous region-of-interest-based time-varying functional connectivity study focusing on the insula, which reported time-varying visual and sensorimotor hyperconnectivity of the ventral aINS. Our findings differ, however, in that we did not detect positive default mode network connectivity to the dorsal aINS and that patterns of hypoconnectivity to primary visual and sensorimotor areas were less apparent in the aforementioned study (15). While left and right aINS subregions primarily differed with respect to connectivity to regions in the ipsilateral hemisphere, consistent differences in time-varying functional connectivity at the system-level were identified when comparing ventral to dorsal aINS subregions. Mirroring static functional connectivity studies and the aforementioned time-varying connectivity study on the insula (7, 8, 15), the task-control Mode 1 of the dorsal aINS showed stronger anti-correlations to the default mode network than did its ventral counterpart. This observation is in line with salience network models proposing that the dorsal aINS receives ventral aINS input streams regarding the moment-to-moment condition of the body and then, based on these inputs, recruits task control (14) and executive control (4) network resources to maintain cognitive set and guide behavior while inhibiting the default mode network (13). The salience network Mode 4 showed stronger connectivity between ventral aINS and ventromedial prefrontal, orbitofrontal, and temporopolar cortices when compared to its dorsal aINS counterpart. This finding is in line with functional-anatomical models suggesting a close alliance between the salience and the semantic appraisal networks (13, 18, 54), whose main components (the temporal pole, ventral striatum, medial orbitofrontal cortex, and amygdala) have been proposed to interact with autonomic representations in the aINS to construct the meaning and significance of social and non-social stimuli under prevailing conditions (13, 18, 54). Importantly, we did not find any differences across aINS subregions in the number of transitions or in the diversity of fractional occupancy profiles. Moreover, our analyses exploring overlapping mode occupancy across aINS subregions revealed that distinct aINS subregions tend to occupy the same mode at the same time. Further support for this conclusion comes from the clustering analysis of individual fractional occupancy profiles, which led to the identification of four meta-profiles characterized by occupancy of a dominant mode. Although our findings are not conclusive on whether aINS subregions always engage the same mode in time, they comprehensively suggest a tendency for coherent mode occupancy that could potentially underpin specific trait characteristics or behavioral states. Future studies could leverage methods such as directed and effective connectivity to assess the temporal relationships between intrinsic time-varying activity of distinct aINS subregions, which could be lagged, phase-shifted, or anticorrelated over time. In order to assess the specificity of aINS time-varying functional connectivity modes, we assessed and compared time-varying functional connectivity of the left IFG, a region involved in language processing and syntax production (43, 52, 55). The left IFG showed region-specific contributions from language processing regions but also displayed pronounced anticorrelation to default mode network regions in Mode 1 and similar time-varying connectivity to the same sensory regions found in Modes 2 and 3 derived from the aINS. These findings suggest that distinct brain regions, although characterized by seed specific connectivity patterns, may cohesively transition between broader modes characterized by shared regional time-varying functional connectivity patterns, possibly revealing a fundamental property of functional brain organization.

### “State” characteristics of aINS time-varying functional connectivity modes

Having determined that two of the four major modes of aINS connectivity involved varying connectivity to sensory, and, most notably, visual cortices, we chose to evaluate how sensory input modifies aINS connectivity dynamics. To this end, we studied the same participants under eyes closed and eyes open scanning conditions. Compared to the eyes open condition, participants with eyes closed spent significantly less time in the task-control network Mode 1 but more time in the primary sensory anti-correlated Mode 2. This finding is consistent with recent static connectivity studies revealing occipital and sensorimotor functional changes in eyes closed versus open scanning conditions (56, 57) and suggests that environmental conditions impact time-varying aINS connectivity by shifting the major modes occupied. Therefore, a measurable component of time-varying connectivity may reflect external cues such as conditions of eyes closure (38, 58), mental states (59, 60), or internally driven physiological conditions of the body (12). Finally, it is tempting to speculate that aINS occupancy of the visually anticorrelated Mode 2 might reflect levels of drowsiness and fluctuations in alertness. Intriguingly, a recent study investigated wakefulness fluctuations as a source of time-varying functional connectivity by combining simultaneously acquired tf-fMRI and EEG data (61). This study revealed progressive whole-brain hypoconnectivity during deeper sleep stages (N2 and N3) when compared to wakefulness, raising the possibility that the increased aINS fractional occupancy in Mode 2 during eyes closed is related to reduced levels of alertness and wakefulness.

### “Trait” characteristics of aINS time-varying functional connectivity modes

To the extent that modes and profiles of time-varying aINS functional connectivity represent traits, individuals should show stability of these features over time. To assess this stability, we studied a longitudinal dataset of cognitively healthy older adults. The four aINS modes were stable over an average of 9 months in this sample, whether assessed by comparing fractional occupancy at the group level via parametric tests or at the individual level via cosine similarity. These findings converge with several recent studies showing that individual topologies in static functional brain organization are highly stable across time and show unique features with promise for use in precision functional mapping of individual human brains (62–65). In line with previous work associating aINS function with social-emotional functions such as empathy (15, 66, 67), we found that the time aINS subregions cohesively spent in the task-control Mode 1 was positively associated with personal distress, while the time aINS subregions spent in the salience Mode 4 was positively associated with the fantasy score from the Interpersonal Reactivity Index. Caution is advised, however, in the interpretation of these findings since the statistical associations were modest and did not survive correction for the four models used to assess these relationships. Nevertheless, the identified associations were specific to the aINS (compared to the left IFG) and survived when removing outliers, adding tentative support for the trait characteristics of aINS time-varying functional connectivity. Nonetheless, the strongest evidence for a trait component to the identified modes remains their reproducibility in individual subjects. The personal distress subscale measures “self-oriented” feelings of personal anxiety and unease in tense interpersonal settings. The level of personal distress was associated with greater time spent in a mode characterized by dorsal cingulo-opercular regions overlapping with the task-control network and anti-correlations to the default mode network, in line with findings associating connectivity of the anterior cingulate cortex and insula with pre-scan anxiety levels, dispositional anxiety, and affective symptoms in mood disorders (4, 20, 68). The fantasy score reflects the capacity of participants to transpose themselves imaginatively into the feelings and actions of fictitious characters in books, movies, and plays. This capacity requires a high level of social contextualization and involves semantic processes typically associated with regions of the semantic appraisal network, such as the temporal lobes and ventro-medial prefrontal areas (13, 18, 54), the same regions contributing to the salience Mode 4. Importantly, fractional occupancies of time-varying functional connectivity modes derived from the left IFG were not significantly associated with measures of dispositional empathy, suggesting a degree of specificity of these findings to the aINS.

### Limitations and future directions

Although time-varying functional connectivity analyses have been of increasing interest to the human brain mapping community, consensus is still lacking about whether time-varying fluctuations in BOLD signal coherence reflect brain physiology or are mainly due to noise stochastically occurring in the scanner between the acquisition of distinct brain volumes (69). One limitation of this work relates to the use of hard clustering techniques such as k-means to extract modes of time-varying functional connectivity, since distinct modes are likely to reflect extremes in time-varying connectivity gradients rather than discrete clusters of topographical connectivity. Therefore, the ideal number of derived time-varying connectivity modes may vary depending on the temporal and spatial resolution of the used dataset, on the investigated anatomical region, and on the specific question of the researcher. We make no claim that the aINS transitions between only four modes, but since we were interested in large-scale time-varying connectivity patterns, we deemed it impractical to use an overly fine-grained mode decomposition by applying higher clustering solutions. Further, the sliding-window approach has been scrutinized because of its moderate reliability (70) and since its output is prone to influence by head movement, sliding-window length, and other methodical choices (71–73), such as global signal regression. In our study, however, we did not find any significant association between time spent by distinct aINS subregions in a specific mode and summary measures of head movement, as exemplified by the regression analyses associating fractional occupancy and dispositional empathy, which were corrected for mean frame-wise head displacement (42, 74). Importantly, our control analyses using the longitudinal dataset preprocessed with global signal regression resulted in the identification of time-varying functional connectivity modes that highly resembled the time-varying modes derived using data without global signal regression (Supplementary results and Figure S6). The only exception was for the primary sensory hyperconnected Mode 3, which was not identified in three out of four aINS subregions, suggesting that this mode is highly modulated by the global signal. A recent report suggested that noise sources (for instance motion and respiration) can have a temporally lagged effect on the BOLD signal, which is greatly reduced by global signal regression (72). However, a recent study investigating time-varying functional connectivity in mice reliably mapped distinct time-varying patterns of tf-fMRI co-activation that occurred at specific phases of global signal fluctuations (75). These findings suggest that the global signal may partially relate to oscillatory cycling of slowly propagating neural activity, as observed in lag threads of propagated tf-fMRI signal in the human brain (76) and transient variation in calcium co-activation patterns in mice (77). Slow oscillation in the global signal have been proposed to coordinate fluctuating periods of brain network topologies characterized by heightened global integration and shifts in arousal (30, 78).

Finally, the limitations in temporal resolution affecting the sliding-window approach may also explain the inability to detect mode-specific fractional occupancy differences between the left and right aINS, although both regions play a distinct role in autonomic outflow and behavior (11, 12). More sophisticated methods that preserve the temporal richness of tf-fMRI data, such as hidden Markov models and Topological Data Analysis (28, 31), may help elucidate lateralized differences in time-varying connectivity of the left and right aINS. However, the detection of static functional connectivity heterogeneity informed by time-varying fractional occupancy profiles and the reproducibility of our findings across distinct preprocessing pipelines, sliding-window lengths, clustering choices, samples, and time points (within individuals)(79) increase confidence in the biological relevance of findings. In the future, our approach could be extended to analyze time-varying functional connectivity of other brain regions, or to assess time-varying fractional occupancy profiles in psychiatric and neurological disorders characterized by aINS dysfunction.

## Abbreviations

aINS: Anterior insula
BOLD: blood oxygen level dependent
IFG: inferior frontal gyrus
tf-fMRI: task-free functional MRI

## Acknowledgements

We thank the participants for their contributions to neuroscience research.

## Data availability

The data that support the findings of this study are available on request from the corresponding author W.W.S. The data are not publicly available due to them containing information that could compromise the privacy of participants in the study.

## Author’s contributions

L.P and W.W.S conceived the study; all authors designed the experiments; L.P implemented the analysis of the data; L.P. and W.W.S. wrote the manuscript; and all the authors edited the manuscript.

