## Supplement for "State and trait characteristics of anterior insula time-varying functional connectivity"

**Supplementary Material and Methods: 1**

- Control analyses on sliding-window length and k-means clustering solution

**Supplementary Results: 4**

- aINS time-varying functional connectivity modes with sliding-window of length 72 TR (144 s)
- aINS time-varying functional connectivity modes with k-means solutions k = 3, 5, and 6
- The impact of global-signal regression on aINS time-varying functional connectivity modes

**Supplementary References**

**Supplementary Figures, Legends and Tables: 15**

- Figure S1: Summary of analyses
- Figure S2: aINS time-varying functional connectivity modes at longer sliding-window length
- Figure S3: aINS time-varying functional connectivity modes with different k-means solutions
- Figure S4: Brain network templates
- Figure S5: Within-group and between-group spatial similarity of aINS modes
- Figure S6: Differences in time-varying connectivity between correspondent modes of the left dorsal and ventral aINS, left and right dorsal aINS and left and right ventral aINS
- Figure S7: Global-signal regression affects aINS time-varying functional connectivity modes
- Figure S8: Transition probability matrices of aINS time-varying connectivity modes
- Figure S9: Silhouette plots for clustering solution k=4 performed on fractional occupancy of aINS subregions
- Figure S10. Static functional connectivity of aINS subregions compared across meta-profiles
- Figure S11. Time-varying functional connectivity modes of the left IFG
- Table S1: Relative contribution of large-scale brain networks to aINS time-varying connectivity modes
- Table S2. Paired t-test statistics for the longitudinal and eyes closed/open datasets
- Table S3. Regression models interpersonal reactivity index subscales
- Video 1: Time-varying functional connectivity maps of the left ventral aINS in an exemplary study participant

**Appendix**

- Code used to generate time-varying functional connectivity maps

**Supplementary Materials and Methods**

*Control analyses on sliding-window length and k-means clustering solution.* To assess the influence of sliding window length on our findings, for the cross-sectional sample, we generated time-varying functional connectivity maps for each aINS subregion using a sliding-window of 72 TR (144 seconds), four times longer than the 18 TR window used to produce the results presented in the main text of the manuscript. For this data, k-means clustering with k = 4 was used to extract time-varying functional connectivity modes of the aINS. Differences in aINS fractional occupancy profiles derived with sliding-window lengths of 18 TR and 72 TR where compared via paired t-tests (p < 0.05 uncorrected). Also on the cross-sectional sample, data generated with a sliding-window length of 18 TR (36 seconds) was further analyzed with k-means clustering solutions of k = 3, 5, and 6. Spatial similarity was assessed between mode maps derived from the same aINS subregion to identify corresponding time-varying functional connectivity modes across distinct k-means clustering solutions. To compute the spatial similarity, Pearson’s correlation coefficients were performed by considering only z-values greater than 0.3 and less than -0.3. With increasing numbers of unassigned modes per higher clustering solution, only the highest correlation coefficient to the mode from the smaller clustering solution was reported.

**Supplementary Results**

*aINS time-varying functional connectivity modes with sliding-window of length 72 TR (144 s).* Spatial maps of time-varying functional connectivity modes of the aINS derived with a sliding-window length of 72 TR (144 seconds) highly resembled the maps derived with sliding-window length of 18 TR (36 seconds) (Supplementary Figure S2). Significant differences in fractional occupancy were not observed for corresponding time-varying aINS modes across the two sliding window lengths, with the one exception that fractional occupancy of the left ventral aINS in the primary sensory anticorrelated Mode 2, showed significantly higher fractional occupancy when derived with a sliding-window length of 72 TRs (p < 0.05 uncorrected). However, the violin plots show that fractional occupancy profiles derived with sliding-window length of 72 TR are not as normally distributed as fractional occupancy profiles derived with sliding-window length of 18 TR. Given the consistency of spatial patterns across sliding-window lengths and the more normal distribution of fractional occupancy when using sliding-window length of 18 TR, we decided to report findings using the shorter sliding-window in the main paper. This decision is further supported by previous studies reporting that shorter sliding-windows are better suited to detect fast transitions in time-varying functional connectivity (1, 2).

*aINS time-varying functional connectivity modes with k-means solutions k = 3, 5, and 6.* Time-varying functional connectivity modes of aINS subregions where derived with clustering solutions of k = 3-6 (Supplementary Figure S3; shown only for left aINS subregions). At lower clustering solutions of k = 3 higher cognitive modes were not detectable as in the k = 4 clustering solution, in particular the salience network Mode 4 was missing for the left dorsal aINS, while the task-control Mode 1 was missing for the left ventral aINS. At higher clustering solutions of k = 5-6, the four modes identified in the k = 4 clustering solution split into redundant sub-modes of aINS time-varying functional connectivity. For the left dorsal aINS, the task-control Mode 1 split into a mode containing only the positive cingulo-opercular correlations, but without anticorrelations to the posterior cingulate and prefrontal cortices (Mode 5, k = 5, spatial correlation to Mode 1 in the lower clustering solution R = 0.61). At even higher clustering solutions, the primary sensory hyperconnected Mode 3 split into a sensorimotor hyperconnected mode (Mode 6, k = 6, spatial correlation to Mode 3 in the lower clustering solution R = 0.94). For the left ventral aINS, the salience Mode 4 split into a mode with stronger contributions from the posterior cingulate (Mode 5, k = 5, spatial correlation to Mode 4 in the lower clustering solution R = 0.74). At even higher clustering solutions, the task-control Mode 1 split into a spatially reduced version, lacking contributions from the dorsal parietal cortex (Mode 6, k = 6, spatial correlation to Mode 1 in the lower clustering solution R = 0.75).

*The impact of global-signal regression on aINS time-varying functional connectivity modes.* K-means clustering of k = 4 was applied on time-varying functional connectivity windows derived from the tf-fMRI longitudinal data sample denoised by including global signal regression. This analysis resulted in four time-varying functional connectivity modes per aINS subregion (Supplementary Figure S7) similar to the time-varying functional connectivity modes derived from the data without global signal regression (Figure 4). All aINS subregions showed time-varying functional connectivity to the task-control Mode 1, to the primary sensory anti-correlated Mode 2, and to the salience Mode 4. Notably, the primary sensory hypercorrelated Mode 3 was identified only in the right ventral aINS, but was not identified in the remaining aINS subregions. Four additional, unidentified modes were generated through this analysis per aINS subregion. In the left dorsal aINS, k-means clustering yielded a mode highly resembling a reduced version of the task-control Mode 1 (Supplementary Figure S7 panel A); in the right dorsal aINS, a mode characterized by general hypoconnectivity was detected (Supplementary Figure S7 panel B); while in the left ventral aINS, k-means clustering yielded a mode resembling a second version of the primary sensory anticorrelated Mode 2 (Supplementary Figure S7 panel C).

**Supplementary Figures, Legends, Tables and Video: 11**

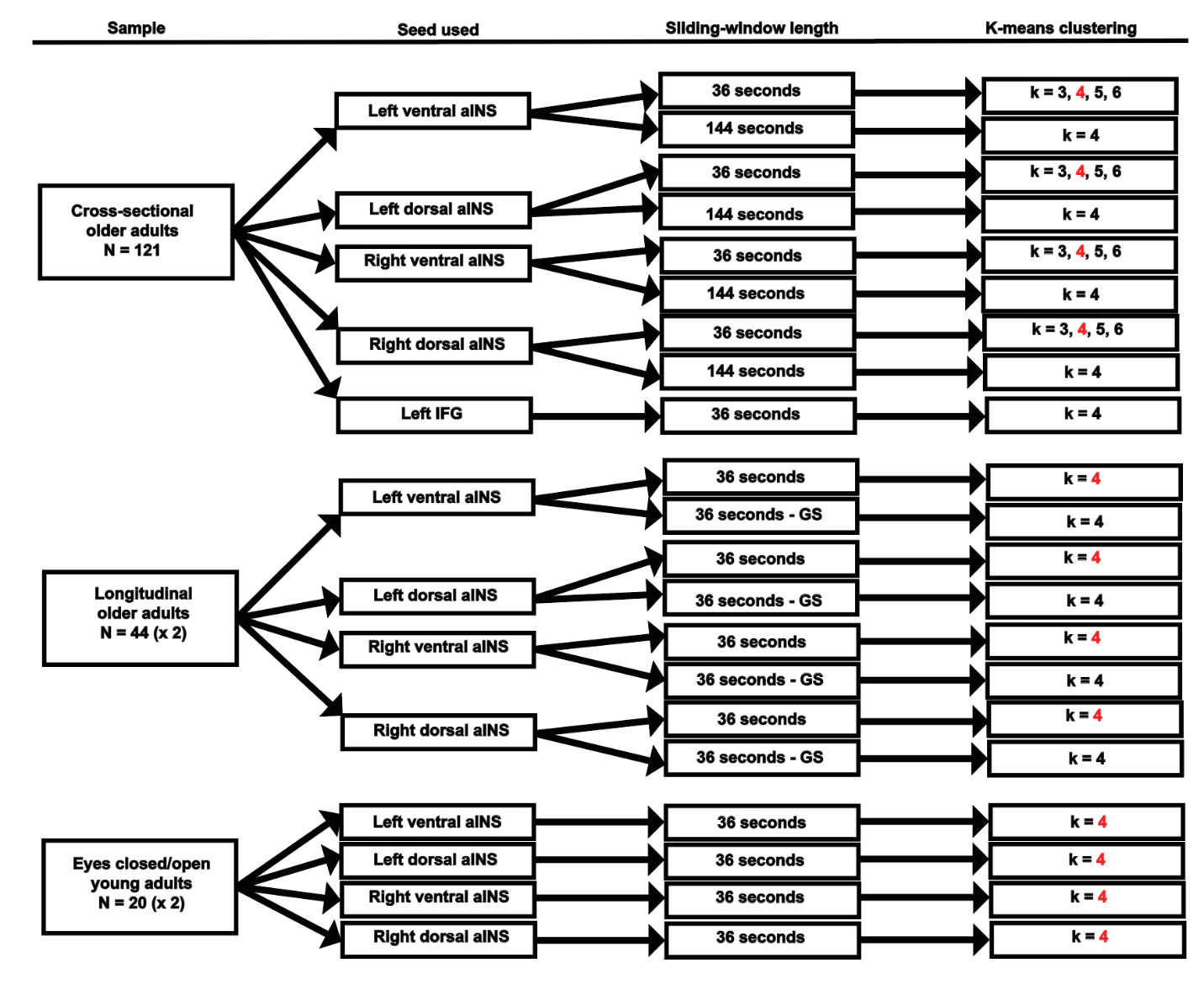

**Supplementary Figure S1. Summary of analyses.** Samples, seeds used, sliding-window length and k-means clustering analyses performed. In red, clustering solutions presented in the main body of the paper. aINS = anterior insula; IFG = inferior frontal gyrus; GS = global signal regressed data.

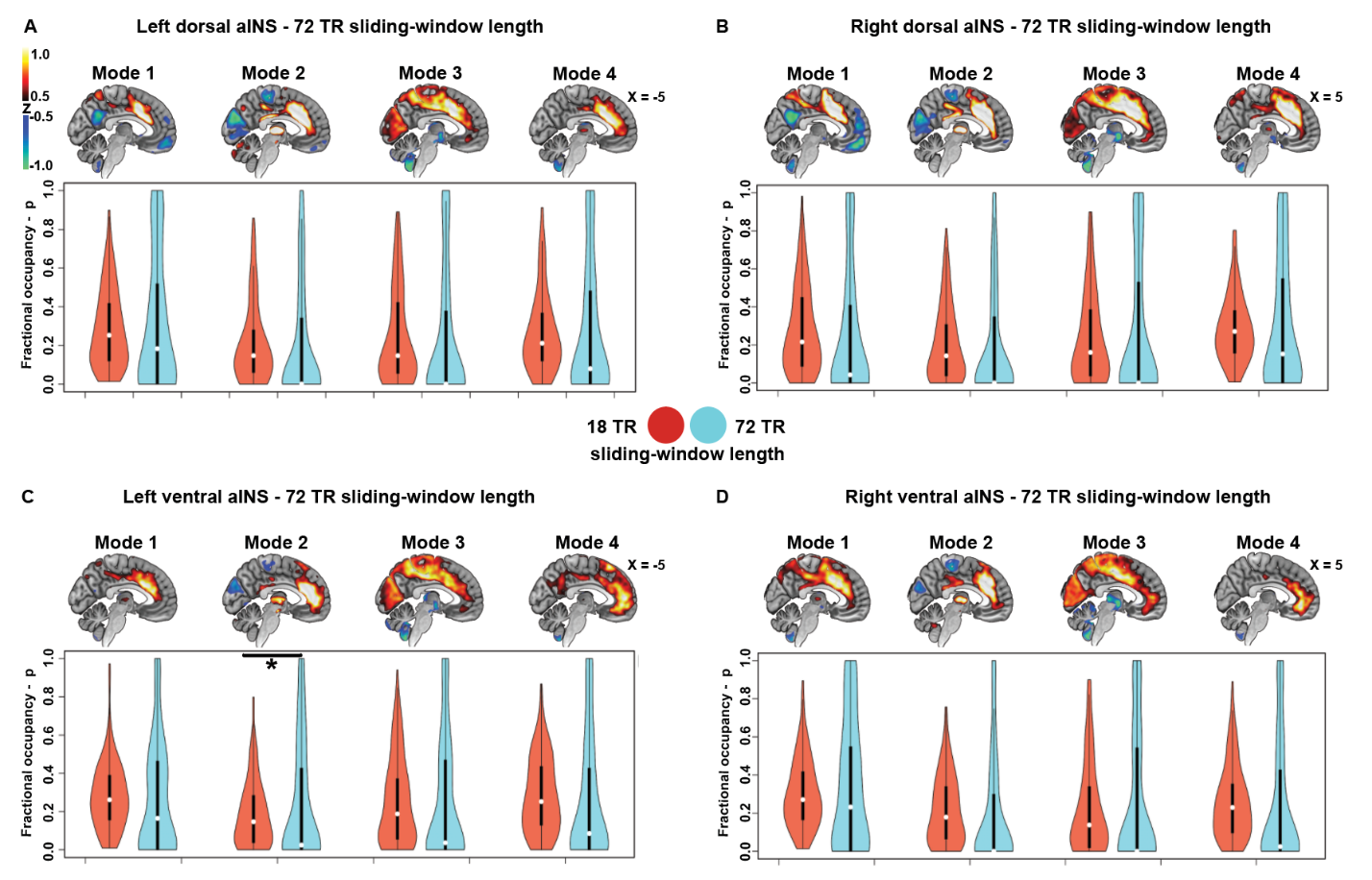

**Supplementary Figure S2. aINS time-varying functional connectivity modes at longer sliding-window length.** In the cross-sectional sample, modes of time-varying functional connectivity derived using a 72 TR sliding-window length showed strong resemblance to those derived with an 18 TR window (sagittal plane only panels **A-D**; threshold at -0.5 > z > 0.5, negative connectivity is depicted in blue, positive in red). Fractional occupancy in corresponding modes did not significantly differ when using sliding-window lengths of 18 TR (red violin plots) versus sliding-window lengths of 72 TR (blue violin plots), with the exception of left ventral aINS fractional occupancy in the primary sensory anticorrelated Mode 2 (paired t-test; *p < 0.05 uncorrected). aINS = anterior insula

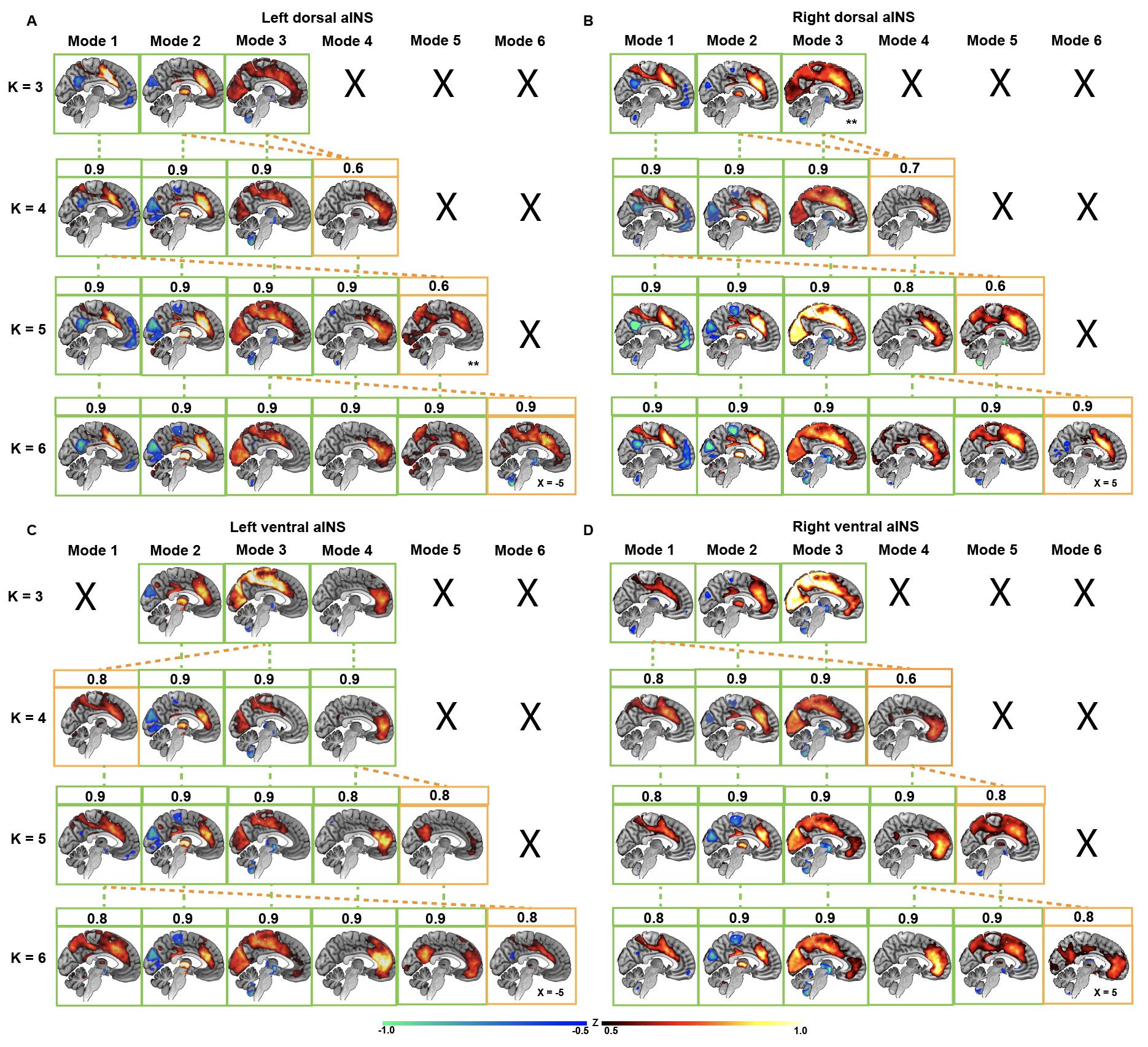

**Supplementary Figure S3. aINS time-varying functional connectivity modes with different k-means solutions.** Time-varying functional connectivity modes of the left dorsal **(A)**, right dorsal **(B)**, left ventral **(C)**, and right ventral **(D)** aINS derived with k-means clustering solutions of k = 3-6. Green connecting lines and squares connect corresponding time-varying functional connectivity modes found across clustering solutions; crosses reflect absence of corresponding modes. Orange connecting lines and square connect new modes that emerge with higher clustering solutions to modes identified with lower-dimensionality reductions. Numbers in boxes reflects highest spatial similarity between higher- and lower-dimensionality modes based on Pearson’s correlation coefficients (multiple corresponding modes possible based on Pearson’s correlation coefficient). aINS = anterior insula; ** mode maps thresholded at values > 0.3 and < -0.3 for visualization purposes

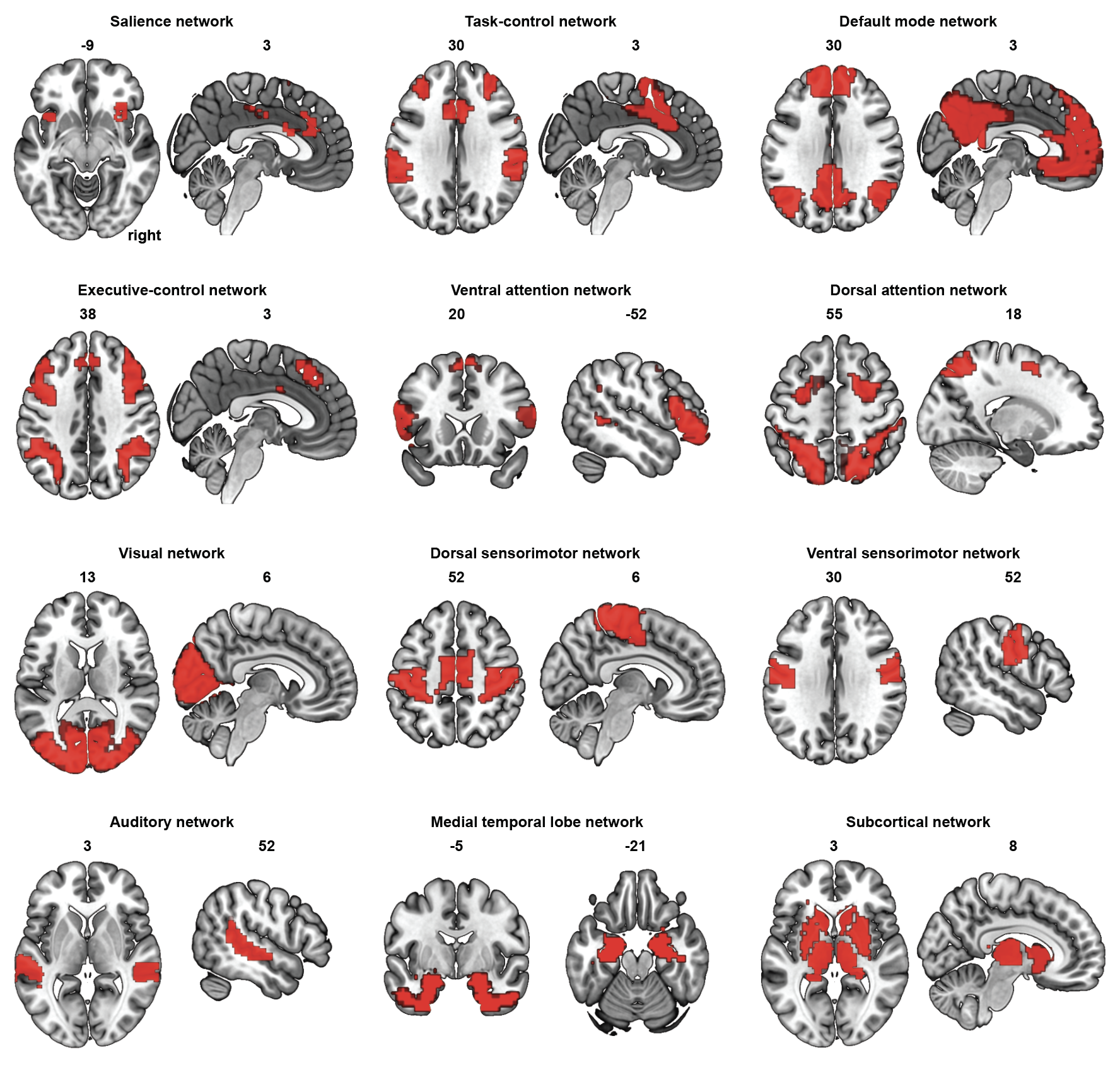

**Supplementary Figure S4**. Publicly available templates of major brain networks derived from a study investigating the functional network organization of the human brain (3) (<https://www.jonathanpower.net/2011-neuron-bigbrain.html>). Subgraphs corresponding to major brain systems were derived from tf-fMRI data of > 300 healthy adults by retaining 2% of the strongest correlations. This procedure resulted in 12 binary templates spanning cognitive, primary sensory and subcortical systems: the salience, cingulo-opercular task-control, default, executive-control, ventral attention, dorsal attention, visual, dorsal sensorimotor, ventral sensorimotor, auditory, medial temporal lobe (“memory retrieval”) and subcortical networks.

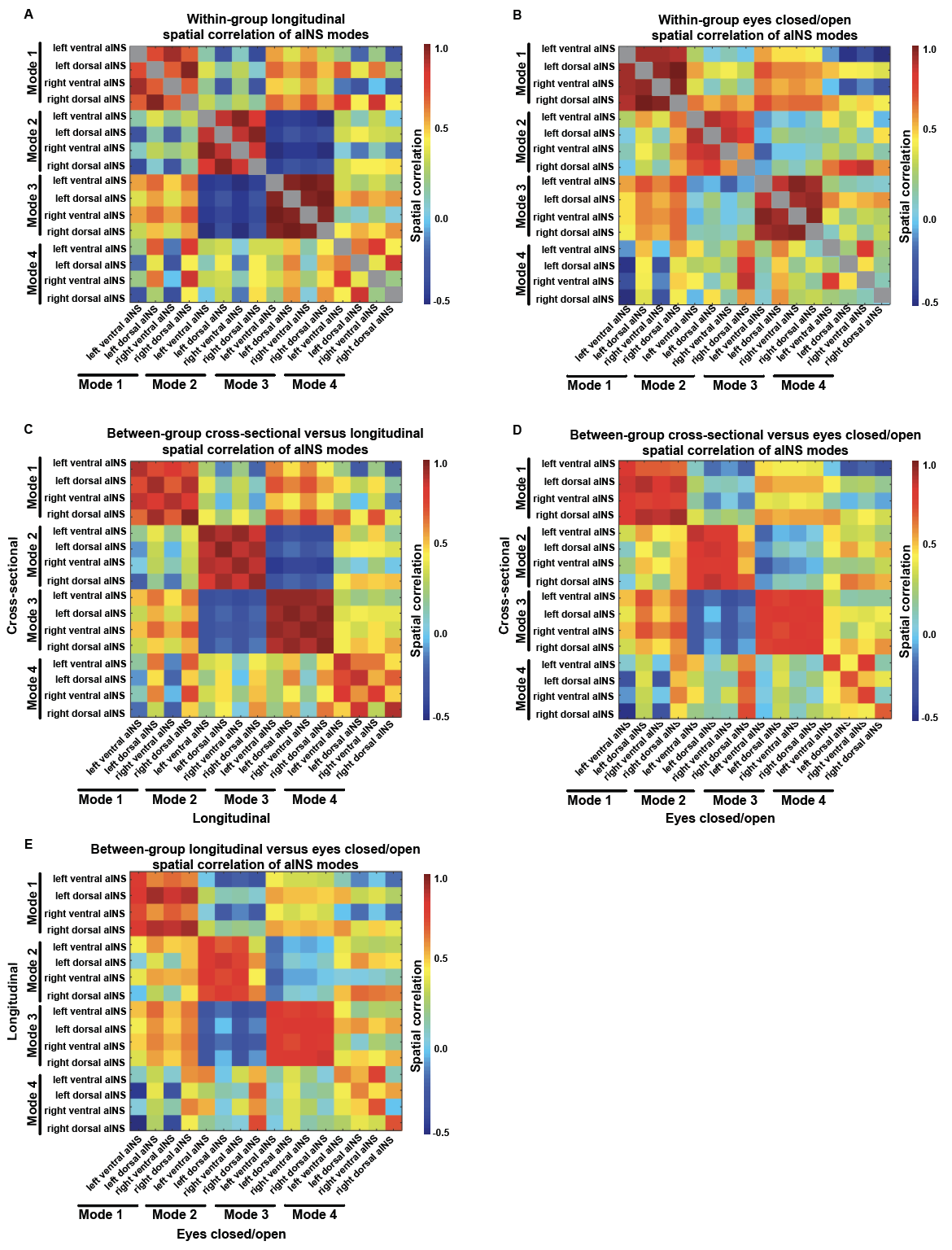

**Supplementary Figure S5.** Within-group and between-group spatial similarity of correspondent aINS modes. **(A)** Spatial similarity of aINS modes within the longitudinal sample. **(B)** Spatial similarity of aINS modes within the eyes closed/open sample. **(C)** Spatial similarity between aINS modes derived from the cross-sectional and longitudinal samples. **(D)** Spatial similarity between aINS modes derived from the cross-sectional and eyes closed/open samples. **(E)** Spatial similarity between aINS modes derived from the longitudinal and eyes closed/open samples. Spatial similarity was assessed via Pearson’s correlation after vectorizing the mode’s templates. Correspondent modes across aINS subregions were characterized by high to moderate levels of spatial similarity both at the within- and between-group levels.

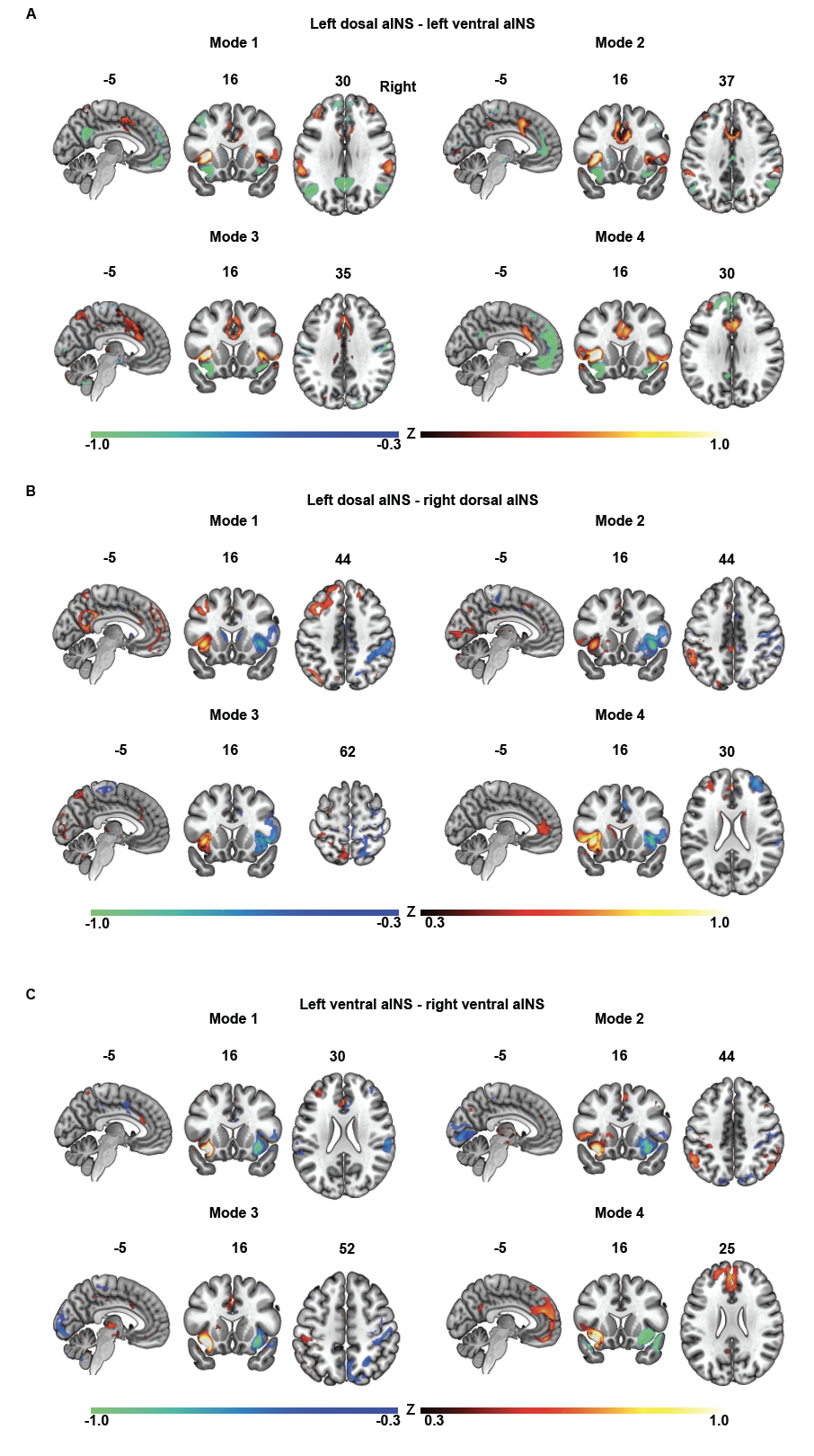

**Supplementary Figure S6.** Differences in time-varying connectivity between correspondent modes of the (**A**) left dorsal and ventral aINS, (**B**) left and right dorsal aINS and (**C**) left and right ventral aINS (threshold at -0.3 > z > 0.3).

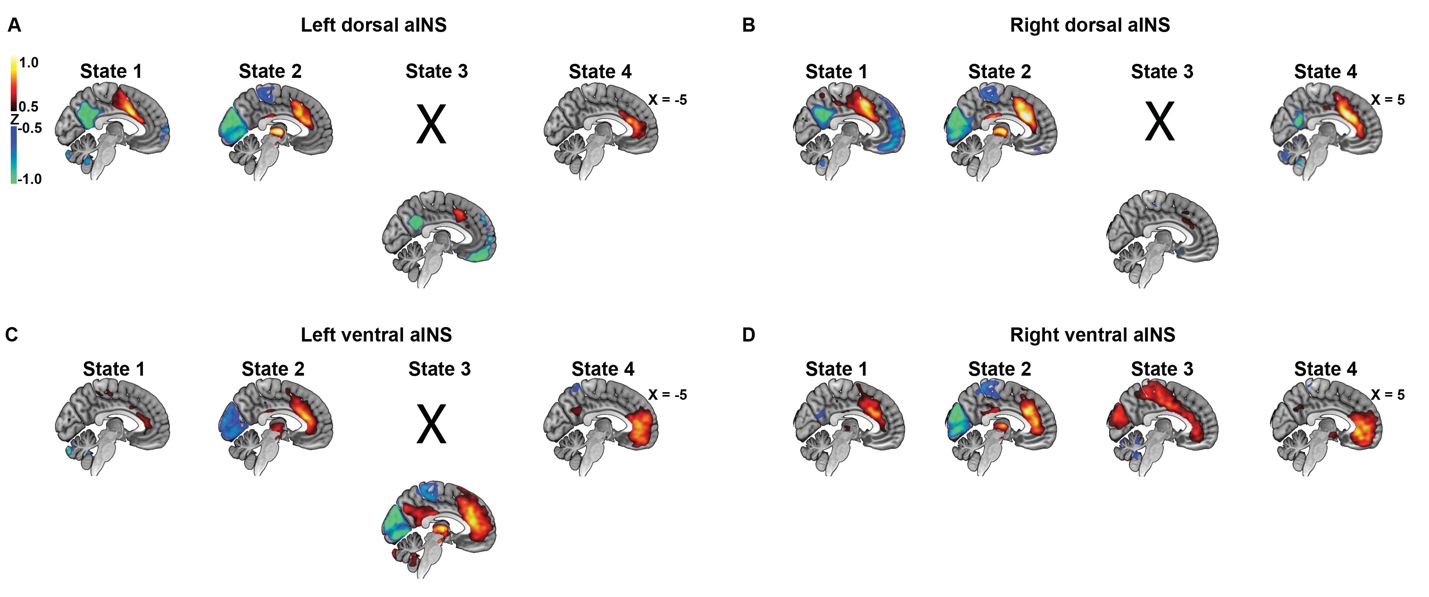

**Supplementary Figure S7. Global-signal regression affects aINS time-varying functional connectivity modes.** In the longitudinal data sample, spatial patterns of time-varying functional connectivity modes where identified across aINS subregions using data preprocessed with global signal regression (sagittal plane only panels **A-D**). The identified time-varying functional connectivity modes show high resemblance to the modes identified in the same data sample without global signal regression. The only exception found was for the primary sensory hypercorrelated mode, which was identified in the right ventral aINS only. K-means clustering instead identified time-varying functional connectivity modes resembling either the task-control Mode 1 (left dorsal aINS, panel A), a general hypocorrelated mode (right dorsal aINS, panel B), or the primary sensory anticorrelated Mode 2 (left ventral aINS, panel C). aINS = anterior insula

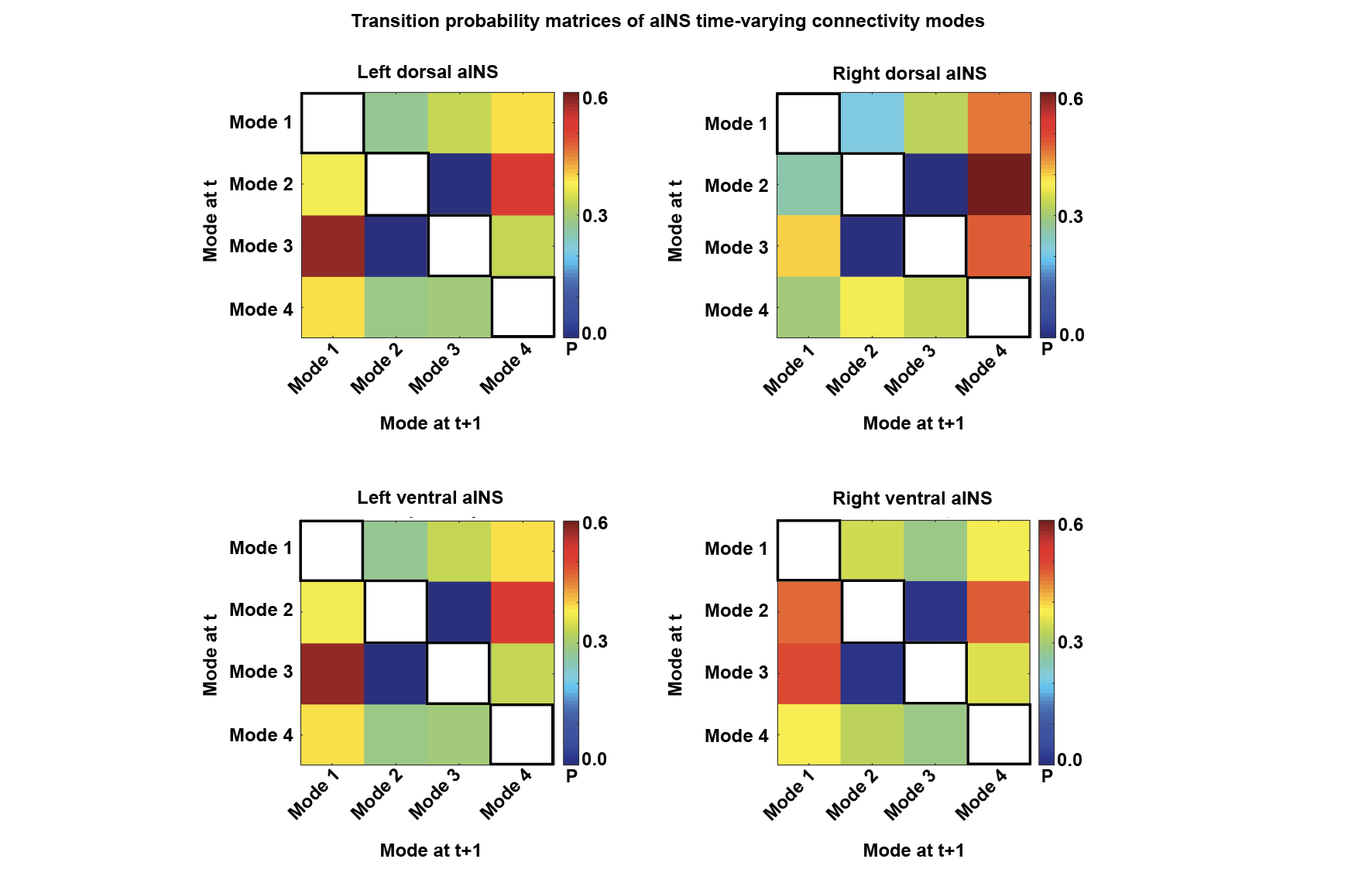

**Supplementary Figure S8.** Transition probability matrices of aINS time-varying connectivity modes. Across aINS subregions, direct transitions between primary sensory anticorrelated Mode 2 and primary sensory correlated Mode 3 did not occur. On-diagonal transitions (from a mode to itself, i.e. dwelling probability) is left blank in order to highlight transition probabilities between distinct modes. Scale bars reflects transition probabilities.

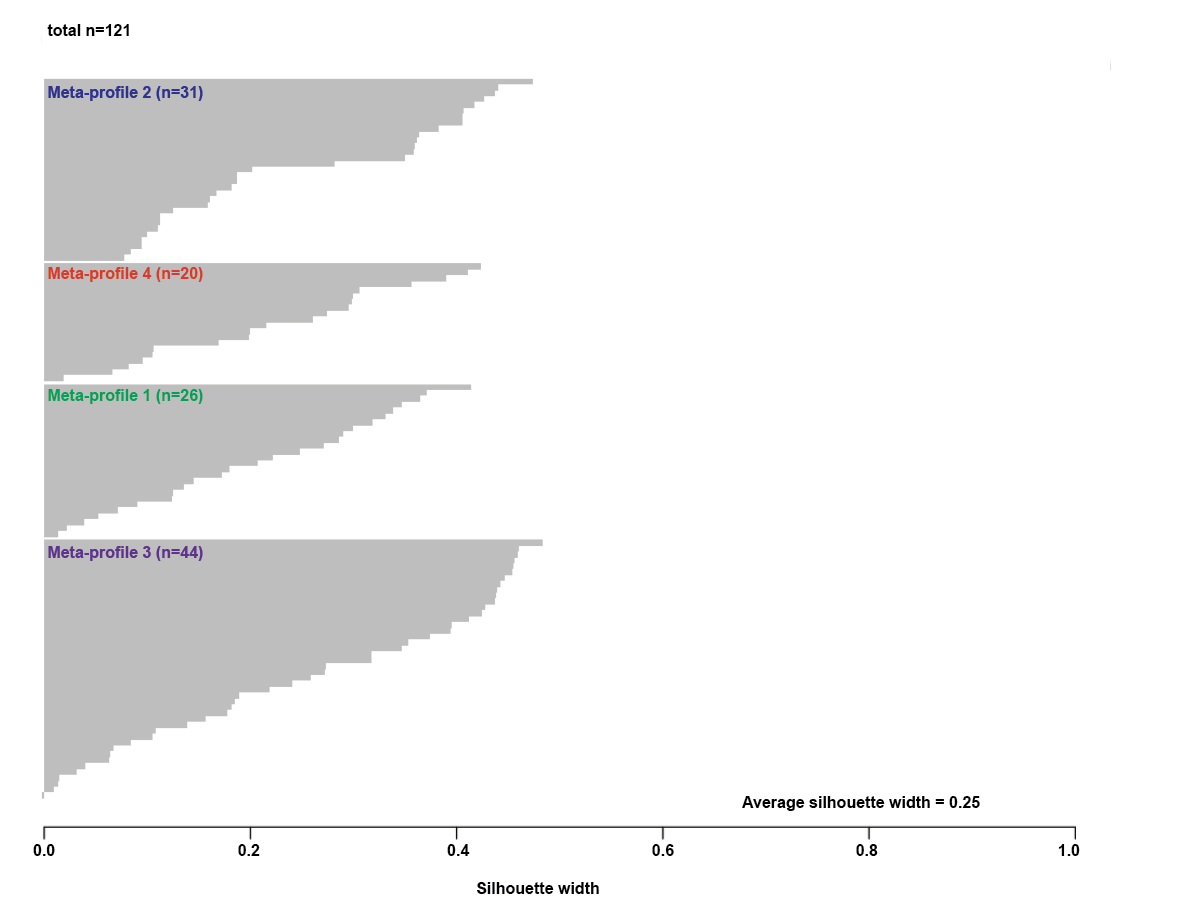

**Supplementary Figure S9.** Silhouette plots for clustering solution k=4 performed on fractional occupancy of aINS subregions.

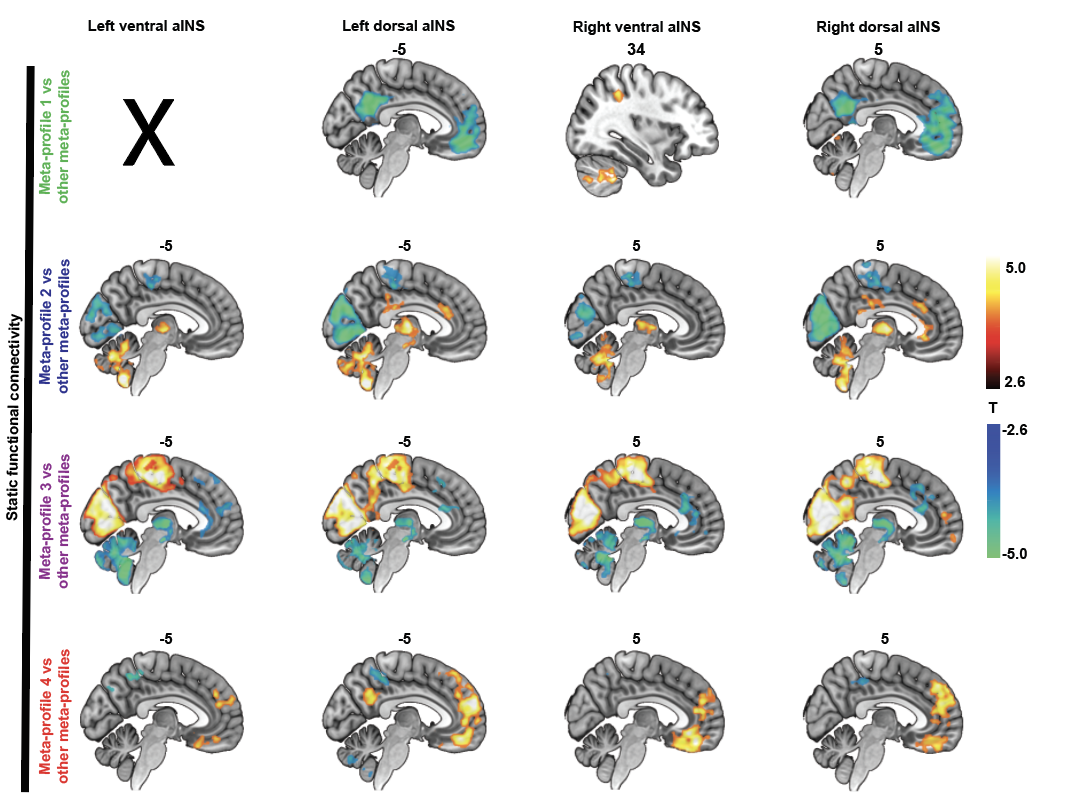

**Supplementary Figure S10.** Static functional connectivity of aINS subregions was compared using ANOVA models and related post-hoc t-tests (height threshold p<0.005; extent threshold p<0.05 FWE-corrected for multiple comparisons), across participants clustered into distinct time-varying functional connectivity meta-profiles. Across aINS subregions, participants in Meta-profile 1 showed prominent static anticorrelation of the dorsal aINS to the default mode network when compared to participants in other meta-profiles; participants in Meta-profiles 2 and 3 showed marked visual static hypoconnectivity and hyperconnectivity; while participants in Meta-profile 4 showed prominent ventral frontal patterns of static connectivity compared to other meta-profiles.

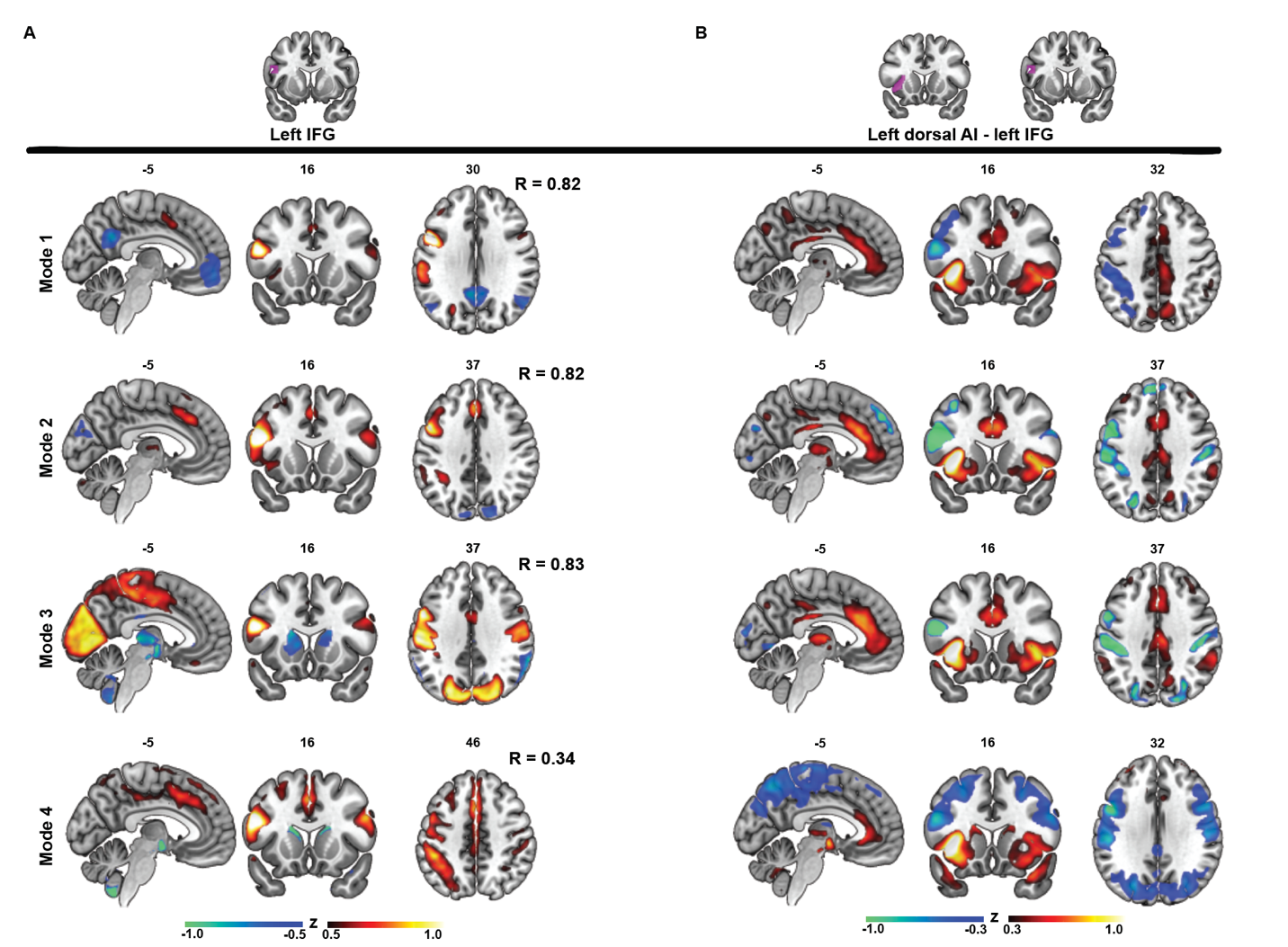

**Supplementary Figure S11.** Time-varying functional connectivity modes of the left IFG. **(A)** Spatial maps of the four time-varying functional connectivity modes identified in the left IFG in the cross-sectional dataset. Spatial similarity to corresponding time-varying functional connectivity modes of the left dorsal aINS was assessed through Pearson’s correlation. Maps thresholded at -0.5 > z > 0.5, negative connectivity is depicted in blue, positive in red. Right hemisphere is on the right side. **(B)** Differences in time-varying connectivity between correspondent modes of the left IFG and left dorsal aINS (threshold at -0.3 > z > 0.3, higher connectivity of the left dorsal aINS is depicted in red, higher connectivity of the left IFG in blue).

|  | **aINS** | **SN** | **TCN** | **DMN** | **ECN** | **VAN** | **DAN** | **VN** | **dSMN** | **vSMN** | **AUN** | **MTL** | **SUB** |
| --- | --- | --- | --- | --- | --- | --- | --- | --- | --- | --- | --- | --- | --- |
| **Mode 1** | left ventral | 0.28 | 0.26 | -0.03 | 0.16 | 0.11 | 0.19 | 0.02 | 0.08 | 0.09 | 0.05 | -0.12 | 0.05 |
|  | right ventral | 0.23 | **0.33** | -0.07 | 0.16 | 0.07 | 0.21 | 0.02 | 0.12 | 0.14 | 0.12 | -0.08 | 0.03 |
|  | left dorsal | 0.18 | **0.39** | **-0.25** | 0.15 | 0.06 | 0.26 | 0.07 | 0.08 | 0.09 | 0.07 | -0.10 | 0.03 |
|  | right dorsal | 0.15 | **0.44** | **-0.33** | 0.15 | 0.00 | **0.30** | 0.09 | 0.15 | 0.16 | 0.10 | -0.10 | 0.02 |
| **Mode 2** | left ventral | **0.47** | 0.14 | 0.20 | 0.23 | 0.26 | **-0.24** | **-0.52** | **-0.37** | **-0.28** | -0.05 | 0.09 | 0.24 |
|  | right ventral | **0.44** | 0.15 | 0.15 | 0.24 | 0.22 | **-0.20** | **-0.38** | **-0.35** | **-0.26** | -0.04 | 0.06 | 0.19 |
|  | left dorsal | **0.51** | 0.28 | 0.08 | **0.31** | 0.24 | -0.14 | **-0.48** | **-0.36** | **-0.28** | -0.08 | -0.03 | 0.23 |
|  | right dorsal | **0.51** | **0.35** | 0.01 | 0.28 | 0.18 | -0.17 | **-0.49** | **-0.32** | **-0.21** | -0.05 | 0.01 | 0.23 |
| **Mode 3** | left ventral | 0.17 | 0.28 | 0.12 | 0.04 | 0.13 | 0.26 | **0.37** | **0.39** | **0.37** | 0.23 | -0.11 | -0.13 |
|  | right ventral | 0.14 | 0.24 | 0.12 | 0.06 | 0.11 | **0.30** | **0.44** | **0.40** | **0.38** | 0.24 | **-0.23** | -0.17 |
|  | left dorsal | 0.26 | **0.38** | 0.07 | 0.16 | 0.15 | **0.31** | **0.34** | **0.35** | **0.34** | 0.20 | **-0.24** | -0.09 |
|  | right dorsal | 0.24 | **0.41** | 0.06 | 0.17 | 0.12 | **0.32** | **0.33** | **0.38** | **0.38** | 0.20 | **-0.27** | -0.11 |
| **Mode 4** | left ventral | **0.30** | 0.08 | 0.27 | -0.01 | 0.23 | **-0.24** | -0.03 | -0.10 | 0.01 | 0.14 | 0.10 | 0.06 |
|  | right ventral | 0.17 | 0.14 | 0.17 | -0.11 | 0.09 | **-0.21** | 0.06 | -0.03 | 0.05 | 0.23 | 0.24 | 0.03 |
|  | left dorsal | **0.33** | **0.30** | 0.10 | 0.06 | 0.28 | -0.17 | 0.02 | -0.13 | 0.03 | 0.20 | 0.06 | 0.07 |
|  | right dorsal | **0.35** | **0.30** | 0.11 | 0.18 | 0.18 | -0.03 | 0.01 | -0.09 | 0.05 | 0.17 | -0.09 | 0.04 |

**Supplementary Table S1.** **Relative contribution of large-scale brain networks to aINS time-varying connectivity modes**. For each aINS subregion, the averaged z-score value enclosed within major brain network templates (3) was extracted from the mode-specific centroid maps generated in the cross-sectional dataset. To appreciate the relative contribution of each large-scale brain network to aINS time-varying connectivity modes, average z-score values ≥ 0.30 or ≤ -0.20 are highlighted in bold font. AUN = auditory network; DAN = dorsal attention network; DMN = default mode network; ECN = executive-control network; MTL = medial temporal lobe “memory retrieval” network; dSMN = dorsal sensorimotor network; vSMN = ventral sensorimotor network; SUB = subcortical network; TCN = (cingulo-opercular) task-control network; VAN = ventral attention network; VN = visual network.

| **Paired t-test statistics: first scanning time point versus second scanning time point** | | | | |
| --- | --- | --- | --- | --- |
|  | **Left ventral aINS** | **Left dorsal aINS** | **Right ventral aINS** | **Right dorsal aINS** |
| **Mode 1** | t = 0.5; p = 0.605 | t = 1.6; p = 0.117 | t = 1.1; p = 0.263 | t = 1.8; p = 0.081 |
| **Mode 2** | t = 0.6; p = 0.540 | t = -1.1; p = 0.289 | t = -0.5; p = 0.600 | t = -1.8; p = 0.075 |
| **Mode 3** | t = -0.6; p = 0.572 | t = -0.1; p = 0.940 | t = -0.4; p = 0.660 | t = 0.0; p = 0.976 |
| **Mode 4** | t = -0.4; p = 0.660 | t = 0.6; p = 0.513 | t = -0.1; p = 0.879 | t = -0.0; p = 0.989 |
| **Paired t-test statistics: eyes closed versus eyes open scanning conditions** | | | | |
|  | **Left ventral aINS** | **Left dorsal aINS** | **Right ventral aINS** | **Right dorsal aINS** |
| **Mode 1** | **t = -3.4; p < 0.003** | **t = -2.1; p < 0.046** | t = -1.9; p = 0.080 | **t = -3.2; p < 0.005** |
| **Mode 2** | **t = 2.3; p < 0.034** | **t = 2.4; p < 0.025** | t = 0.1; p = 0.889 | **t = 3.0; p < 0.007** |
| **Mode 3** | t = 1.2; p = 0.225 | t = 0.5; p = 0.619 | t = 0.3; p = 0.749 | t = 0.3; p = 0.788 |
| **Mode 4** | t = 0.6; p = 0.581 | t = -0.1; p = 0.898 | t = 0.8; p = 0.426 | t = 0.2; p = 0.851 |

**Supplementary Table S2. Paired t-test statistics**, comparing the time spent by aINS subregions in the four time-varying functional connectivity modes in longitudinal dataset and the eyes closed vs eyes open datasets. Significant differences (p < 0.05 uncorrected for multiple comparisons) highlighted in bold. aINS = anterior insula

| **Regression coefficients** | | | | | | | |
| --- | --- | --- | --- | --- | --- | --- | --- |
| **Empathic concern** | **Mode 1** | **Mode 2** | **Mode 3** | **Mode 4** | **Age** | **Sex** | **Sum FWHD** |
| **Model 1: Average aINS** | 0.0 | -3.6 | -3.9 | -1.2 | 0.0 | **3.0*** | 0.0 |
| **Model 2: Left IFG** | -54.7 | -52.7 | -54.7 | -54 | 0.0 | **2.8*** | 0.0 |
| **Perspective** **taking** | **Mode 1** | **Mode 2** | **Mode 3** | **Mode 4** | **Age** | **Sex** | **Sum FWHD** |
| **Model 3: Average aINS** | -0.8 | 1.1 | -5.3 | 0.9 | -0.1 | 1.1 | 0.0 |
| **Model 4: Left IFG** | -57.7 | -58.0 | -60.3 | -61.1 | -0.1 | 1.3 | 0.0 |
| **Personal distress** | **Mode 1** | **Mode 2** | **Mode 3** | **Mode 4** | **Age** | **Sex** | **Sum FWHD** |
| **Model 5: Average aINS** | **7.1*** | -2.3 | -3.0 | -1.6 | 0.0 | 0.8 | 0.0 |
| **Model 6: Left IFG** | -4.2 | -5.1 | 0.2 | -5.3 | 0.0 | 1.0 | 0.0 |
| **Fantasy score** | **Mode 1** | **Mode 2** | **Mode 3** | **Mode 4** | **Age** | **Sex** | **Sum FWHD** |
| **Model 7: Average aINS** | -1.2 | 3.2 | 1.1 | **9.2*** | 0.1 | 1.8 | 0.0 |
| **Model 8: Left IFG** | -52.9 | -56.7 | -57.6 | -56.9 | 0.1 | **2.3*** | 0.0 |

**Supplementary Table S3. Regression coefficients and significance level** for the eight multiple linear regression models using subscales of the Interpersonal Reactivity Index as dependent variables and associating these subscales with time spent in the four modes, once averaged across aINS subregions, once for the left IFG. * p < 0.03 uncorrected for multiple comparisons, highlighted in bold. aINS = anterior insula, FWHD = frame-wise head displacement, IFG = inferior frontal gyrus

**Video 1.** Lateral and mid-sagittal views of 218 concatenated time-varying functional connectivity maps of the left ventral aINS in a typical study participant. Red reflects positive, blue negative connectivity to the left ventral aINS. Only z-values of time-varying functional connectivity higher than 1.2 and smaller than -1.2 are shown. The right hemisphere is shown on the right side of the video.

**Appendix**

Code used to generate time-varying functional connectivity maps. The code was implemented in Python and leverages FSL-based functions. The code is also available in GitLab (<https://gitlab.com/juglans/sliding-window-analysis/tree/master>)

1. **import** argparse
2. **from** nipype **import** Node, Workflow, Function
3. **from** nipype.interfaces.utility **import** Merge as niMerge
4. **from** nipype.interfaces.fsl **import** Merge as FSLMerge
5. **from** nipype.interfaces.fsl.utils **import** ExtractROI
6. **from** nipype.interfaces.fsl **import** ImageMeants, GLM
7. **from** nibabel **import** load, Nifti1Image
8. **import** numpy as np
9. **import** os
10. **from** sys **import** exit
12. **def** parse_args():
13. parser = argparse.ArgumentParser()
15. parser.add_argument('input_flist', help='Path of input file list, must be newline delimited text file with one file path to a .nii or .nii.gz file on each line.')
16. parser.add_argument('seed', help='Path to the desired seed mask image. If the image is not binary it will be assumed that all nonzero voxels in the mask image are desired for timeseries extraction.')
17. parser.add_argument('--output_path', help='Path to desired output directory. If the path does not exist it will be created.')
18. parser.add_argument('--slidingstep', type=int, default=1, help='Number of TRs to increment the sliding windows.')
19. parser.add_argument('--slidingwindow', type=int, default=18, help='Number of TRs to include in each sliding window constituent analysis.')
20. parser.add_argument('--tstat', action='store_true', default=False, help='Output and concatenate t-statistics from sliding window analysis.')
21. parser.add_argument('--zstat', action='store_true', default=True, help='Output and concatenate z-statistics from sliding window analysis.')
22. parser.add_argument('--bstat', action='store_true', default=False, help='Output and concatenate betas (parameter estimates) from sliding window analysis.')
23. parser.add_argument('--workingdir', type=str, default='/tmp', help='Working directory to store temp nipype workflow files and intermediates. Defaults to /tmp or TMPDIR (if created).')
24. parser.add_argument('--n-procs', type=int, default=2, help='Number of processes (not threads) to use for the MultiProcessing.')
26. args = parser.parse_args()
28. **if** os.path.exists(args.input_flist) **is** False:
29. **print**('Input file [{}] does not exist. Exiting.'.format(args.input_flist))
30. exit(1)
32. **if** **not** args.output_path:
33. **print**('Output directory path was not specified, writing outputs to ./sliding_window_analysis/')
34. args.output_path = os.path.join(os.getcwd(), 'sliding_window_analysis')
35. **else**:
36. args.output_path = os.path.abspath(args.output_path)
38. **if** os.path.isdir(args.output_path) **is** False:
39. **print**('Creating output directory: {}'.format(args.output_path))
40. os.makedirs(args.output_path, exist_ok=True)


44. **return** args
46. **def** isnii(nii_file):
47. **try**:
48. load(nii_file)
49. **return** True
50. **except** TypeError:
51. **print**('Could not open {} using nibabel. Check for file integrity? This file will be skipped.')
52. **return** False
53. **except** FileNotFoundError:
54. **print**('Could not find {}. This file will be skipped.')
55. **return** False
57. **def** get_binarized_seed(seedpath):
58. **print**('Checking if seed is binarized.')
59. nii_obj  = load(seedpath)
60. nii_data = nii_obj.get_fdata()
62. num_zero_or_one_vox = np.count_nonzero((nii_data == 0) | (nii_data == 1))
63. **if** num_zero_or_one_vox == nii_data.size:
64. **print**('Seed is binary.')
65. **return** seedpath
66. **else**:
67. binarized = nii_data.copy()
68. binarized[binarized > 0] = 1
70. seed_dirname, seed_basename = os.path.split(seedpath)
71. fstem = seed_basename.split('.')[0]
72. binarized_seed_basename = seed_basename + '_binarized.nii.gz'
73. binarized_seed_fullpath = os.path.join(seed_dirname, binarized_seed_basename)
75. **print**('Seed is not binary. Converting to binary nii.gz file @ {}'.format(binarized_seed_fullpath))
77. Nifti1Image(binarized, nii_obj.affine, nii_obj.header).to_filename(binarized_seed_fullpath)
79. **print**('Finished writing binarized seed file.')
81. **return** binarized_seed_fullpath
83. **def** merge_save(nii_paths, output_path):
84. **from** nibabel **import** concat_images
85. concatenated_image = concat_images(nii_paths)
86. concatenated_image.to_filename(output_path)
87. **return** output_path
89. **def** parse_input_files(input_flist):
90. nii_paths = []
91. with open(input_flist, 'r') as f:
92. **for** lineno, line **in** enumerate(f):
93. stripped_line = line.strip()
94. **if** isnii(stripped_line) **is** False:
95. **pass**
96. **else**:
97. nii_paths.append(stripped_line)
98. **if** len(nii_paths) == 0:
99. **print**('No valid nii files were located. Exiting.')
100. exit(1)
102. **print**('Parsed input file. {} lines were counted, and {} filepaths were extracted.'.format(lineno, len(nii_paths)))
103. **return** nii_paths
105. **def** get_sliding_window_indices(total_num_vols, window_step, window_size):
106. window_start_indices = []
107. **for** idx **in** range(0, total_num_vols, window_step):
108. **if** idx + window_size > total_num_vols:
109. **break**
110. window_start_indices.append(idx)
111. **return** window_start_indices, [window_size]*len(window_start_indices)
113. **def** generate_sliding_window_map(nii_file, seed, slidingstep, slidingwindow, base_dir, output_dir, betas=False, tstat=False, zstat=True, identifier=None, seedname=None):
115. wkflow_name = os.path.basename(nii_file).split('.')[0]
116. nii_stem = wkflow_name
117. node_prefix = wkflow_name
118. **if** identifier **is** **not** None:
119. identifier = '_' + identifier
120. wkflow_name = wkflow_name + '_' + identifier
121. **else**: identifier = ''
123. **if** seedname **is** None:
124. seedname = os.path.basename(seed)
126. workflow = Workflow(name=os.path.basename(nii_file).split('.')[0], base_dir=base_dir)
128. nii_obj = load(nii_file)
129. x, y, z, numvols = nii_obj.shape
130. tr = nii_obj.header['pixdim'][3]
132. sliding_start_indices, window_lengths = get_sliding_window_indices(numvols, slidingstep, slidingwindow)
133. num_windows = len(sliding_start_indices)
134. **print**(num_windows)
135. **print**(sliding_start_indices, window_lengths)
137. **if** betas:
138. betas_merged_file = os.path.join(output_dir, nii_stem + '_betas_dynamic-step_{}-window_{}_{}'.format(slidingstep, slidingwindow, seedname))
139. betas_nipype_merge_node = Node(niMerge(num_windows), name=node_prefix + '_BetaWindowCollectorNode' + identifier)
140. betas_file_merge_node = Node(Function(function=merge_save, input_names=['nii_paths', 'output_path'], output_names=['output_path']), output_path=betas_merged_file, name=node_prefix + '_betasFinalMergeNode' + identifier)
141. betas_file_merge_node.inputs.output_path = betas_merged_file
143. **if** tstat:
144. tstat_merged_file = os.path.join(output_dir, nii_stem + '_tstat_dynamic-step_{}-window_{}_{}'.format(slidingstep, slidingwindow, seedname))
145. tstat_nipype_merge_node = Node(niMerge(num_windows), name=node_prefix + '_tstatWindowCollectorNode' + identifier)
146. tstat_file_merge_node = Node(Function(function=merge_save, input_names=['nii_paths', 'output_path'], output_names=['output_path']), output_path=tstat_merged_file, name=node_prefix + '_tstatFinalMergeNode' + identifier)
147. tstat_file_merge_node.inputs.output_path = tstat_merged_file
149. **if** zstat:
150. zstat_merged_file = os.path.join(output_dir, nii_stem + '_zstat_dynamic-step_{}-window_{}_{}.nii.gz'.format(slidingstep, slidingwindow, seedname))
151. zstat_nipype_merge_node = Node(niMerge(num_windows), name=node_prefix + '_zstatWindowCollectorNode' + identifier)
152. zstat_file_merge_node = Node(Function(function=merge_save, input_names=['nii_paths', 'output_path'], output_names=['output_path']), output_path=zstat_merged_file, name=node_prefix + '_zstatFinalMergeNode' + identifier)
153. zstat_file_merge_node.inputs.output_path = zstat_merged_file
155. **for** sliding_window_index, windowparams **in** enumerate(zip(sliding_start_indices, window_lengths)):
156. window_start, window_length = windowparams
158. out_beta = nii_stem + '_window_{}_beta.nii.gz'.format(str(sliding_window_index).zfill(3))
159. out_z = nii_stem + '_window_{}_zstat.nii.gz'.format(str(sliding_window_index).zfill(3))
160. out_t = nii_stem + '_window_{}_tstat.nii.gz'.format(str(sliding_window_index).zfill(3))
162. slicer_node = Node(ExtractROI(in_file=nii_file, t_min=window_start, t_size=window_length), name=node_prefix + '_SlicerNum{}'.format(str(sliding_window_index).zfill(3) + identifier))
163. sliding_ts_extract_node = Node(ImageMeants(mask=seed), name=node_prefix + '_TSExtractNum{}'.format(str(sliding_window_index).zfill(3) + identifier))
164. sliding_glm_node = Node(GLM(out_file=out_beta, out_z_name=out_z, out_t_name=out_t), name=node_prefix + '_GLMNum{}'.format(str(sliding_window_index).zfill(3) + identifier))
166. workflow.connect([
167. (slicer_node, sliding_ts_extract_node, [('roi_file', 'in_file')]),
168. (slicer_node, sliding_glm_node, [('roi_file', 'in_file')]),
169. (sliding_ts_extract_node, sliding_glm_node, [('out_file', 'design')]),
170. ])
172. **if** betas:
173. workflow.connect([(sliding_glm_node, betas_nipype_merge_node, [('out_file', 'in{}'.format(sliding_window_index+1))])])
174. **if** tstat:
175. workflow.connect([(sliding_glm_node, tstat_nipype_merge_node, [('out_t', 'in{}'.format(sliding_window_index+1))])])
176. **if** zstat:
177. workflow.connect([(sliding_glm_node, zstat_nipype_merge_node, [('out_z', 'in{}'.format(sliding_window_index+1))])])
179. **if** betas:
180. workflow.connect([(betas_nipype_merge_node, betas_file_merge_node, [('out', 'nii_paths')])])
181. **if** tstat:
182. workflow.connect([(tstat_nipype_merge_node, tstat_file_merge_node, [('out', 'nii_paths')])])
183. **if** zstat:
184. workflow.connect([(zstat_nipype_merge_node, zstat_file_merge_node, [('out', 'nii_paths')])])
186. **return** workflow
188. **def** sliding_window_analysis(input_flist, output_path, seed, slidingstep, slidingwindow, betas=False, tstat=False, zstat=True, working_dir=None, n_procs=2):
189. nii_paths = parse_input_files(input_flist)
190. binarized_seed = get_binarized_seed(seed)
192. **if** working_dir **is** None:
193. working_dir = '/tmp'
195. workflow = Workflow(name='sliding_window_analysis', base_dir=working_dir)
197. **for** file_idx, nii_file **in** enumerate(nii_paths):
198. single_analysis = generate_sliding_window_map(nii_file, seed, slidingstep, slidingwindow, working_dir, output_path, identifier=str(file_idx).zfill(4), tstat=args.tstat, betas=args.bstat, zstat=args.zstat)
199. workflow.add_nodes([single_analysis])
201. workflow.run(plugin='MultiProc', plugin_args={
202. 'n_procs': args.n_procs,
203. 'crashfile_format': 'txt'})
205. **if** __name__ == '__main__':
206. args = parse_args()
207. sliding_window_analysis(args.input_flist, args.out
